# Fine-tuning of Fgf8 morphogen gradient by heparan sulfate proteoglycans in the extracellular matrix

**DOI:** 10.1101/2023.11.02.565243

**Authors:** Mansi Gupta, Thomas Kurth, Fabian Heinemann, Petra Schwille, Sebastian Keil, Franziska Knopf, Michael Brand

## Abstract

Embryonic development is orchestrated by the action of morphogens, which spread out from a local source and activate, in a field of target cells, different cellular programs based on their concentration gradient. Fibroblast growth factor 8 (Fgf8) is a morphogen with important functions in embryonic organizing centers. It forms a gradient in the extracellular space by free diffusion, interaction with the extracellular matrix (ECM) and receptor-mediated endocytosis. However, morphogen gradient regulation by ECM is still poorly understood. Here we show that specific Heparan Sulfate Proteoglycans (HSPGs) bind Fgf8 directly in the ECM of living zebrafish embryos, thus affecting its diffusion and signaling. Using single-molecule Fluorescence Correlation Spectroscopy, we quantify this binding *in vivo*, and find two different modes of interaction. First, reducing or increasing the concentration of specific HSPGs in the extracellular space alters Fgf8 diffusion, and thus, its gradient shape. Second, ternary complex formation of Fgf8 ligand with Fgf-receptors and HSPGs at the cell surface requires HSPG attachment to the cell membrane. Together, our results show that graded Fgf8 morphogen distribution is achieved by constraining free Fgf8 diffusion through successive interactions with HSPGs at the cell surface and in ECM space.

**Statement of significance:** Fgf8 is a secreted morphogen signaling molecule that instrúcts neighboring arrays of undifferentiated cells about their position and cellular identity in tissue. Fgf8 and other morphogens are often distributed in a graded fashion, and can typically work at very low concentrations. To reproducibly generate information in developing tissue, mechanisms have evolved to carefully control distribution and concentration of Fgf8 morphogen. We show that freely diffusing Fgf8 morphogen moves through interstitial cell spaces on its way to target cells, and while doing so, interacts with ECM molecules in these spaces and at cell surfaces via low affinity, reversible binding. These interactions are important tuning mechanisms that contribute to forming the Fgf8 morphogen gradient and to cell surface receptor binding, and thus, to controlling cell type identity.

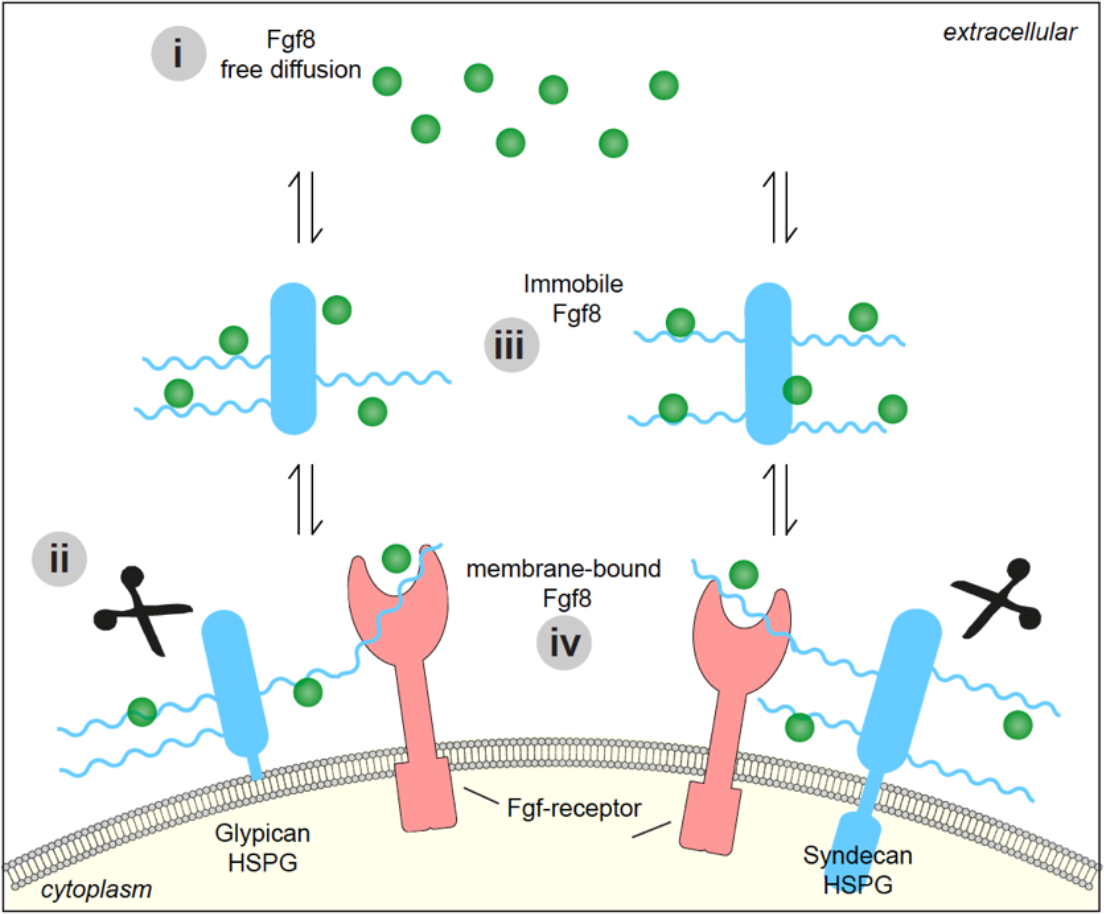

**Graphical abstract:** To generate a gradient, the morphogen Fgf8 shuttles between fast free diffusion through extracellular space (i), slow diffusion or immobility (ii, iii) when bound to Heparan Sulfate Proteoglycans (HSPGs), in the extracellular matrix (ECM) and at cell surface receptors (iv), as revealed by single molecule studies in living zebrafish embryos.

## Introduction

During tissue development, secreted morphogens spread out from a local source through arrays of target cells in a concentration gradient, and activate cellular programs via binding to cell surface receptors to determine cell fate in a concentration dependent manner [1-7]. Fibroblast growth factor 8 (Fgf8) is a morphogen with important functions in several embryonic organizing centers and inductive interactions [8-11]. By freely diffusing through extracellular spaces, Fgf8 forms a concentration gradient that is thought to be shaped by interaction with the extracellular matrix (ECM) and receptor-mediated endocytosis [3, 12, 13],. During early gastrulation, the majority of Fgf8 molecules (91 %) spread in between cells by free diffusion, as was seen using Fluorescence Correlation Spectroscopy (FCS) in living zebrafish embryos, but a small fraction of the molecules (9 %) move as a slow fraction, which is dependent on extracellular heparan sulfate (HS) sugar chains [3, 7]. Novel biophysical approaches to directly monitor transport at the single-molecule level have helped in quantitatively elucidating the mechanisms of transport [3, 6, 7, 14-20], but the exact interplay between free diffusion and interactions with the ECM is not clear, and the identity of molecules in the ECM which regulate Fgf8 transport *in vivo* is not well understood.

Fgf proteins strongly bind to HS sugar chains, which can function as co-receptors for Fgf signal transduction, act as reservoir in the ECM and can also mediate short range mobility of Fgfs along their chains [21-24]. Endogenously, HS chains are attached to core proteins, together constituting Heparan Sulfate Proteoglycans (HSPGs). There are two cell-surface HSPG protein families, namely glypicans (Gpc) and syndecans (Sdc), which differ in their mode of membrane attachment. Glypicans contain a GPI-anchor domain whereas syndecans are connected via a transmembrane domain [25, 26]. Several of these proteins are ubiquitously expressed during early development and can be cleaved from the cell surface into the ECM [27-29]. Individual core proteins can specifically help in the reception of signals like Wnt and Hedgehog [30, 31]. Here, we present a systematic and quantitative analysis of core-protein function with regard to Fgf8 signaling. In particular, our work emphasizes how the different HSPGs influence gradient formation of Fgf8 in the context of living tissues, measured at the single molecule level.

## Results and Discussion

To identify potential HSPG molecules influencing Fgf8 gradient formation, we first examined Fgf8 signal transduction and binding with Gpc and Sdc HSPGs in the presence or absence of their membrane anchorage. We analyzed the expression pattern of Fgf downstream targets *sprouty2* (*spry2*), *sprouty4* (*spry4*) and *ETS translocation variant 4 (etv4)*, under conditions of ectopic Gpc and Sdc expression (Figure S1, Figure S2). In wild-type embryos, *spry2* and *spry4* expression is confined to the margin and *etv4* is expressed more broadly [32-34]. mRNA injection of each full-length Gpc and Sdc HSPG (named Gpc-fl/Sdc-fl), did not influence the expression pattern of *spry2* and *etv4* (Figure S1c,d). In order to examine if cell surface shedding of HSPGs occurs, we generated full-length fluorescent protein fusion constructs, for Gpc4, Gpc3 and Sdc2. Confocal imaging revealed that the fusion proteins are localized on the membrane as well as in the ECM (Figure S2a-c). This suggests that cell-surface shedding is a common mechanism among HSPGs, and raised the possibility of distinct functions of cell-surface bound *versus* extracellular HSPGs. Hence we generated HSPGs lacking membrane anchoring ability, which are confined to the ECM, by replacing their membrane anchors with mRFP, named Gpc^ex^mRFP and Sdc^ex^mRFP (Figure S1a,b). Injection of extracellular Sdc (Sdc2^ex^mRFP, Sdc3^ex^mRFP and Sdc4^ex^mRFP) and specific extracellular Gpc (Gpc1b^ex^mRFP, Gpc4^ex^mRFP and Gpc6a^ex^mRFP) resulted in an expansion of *spry2/spry4* and *etv4* expression domains (Figure S1e-f), reflecting an overall increased range of Fgf8 signal transduction. Interestingly, these three Gpc belong to the same sub-family and all the mentioned Gpc and Sdc are expressed ubiquitously during embryonic development [27, 35]. An alternative possibility of increased Fgf8 production at the source was ruled out, since *fgf8* mRNA expression itself remained unaltered (Figure S1g). Also, control injection of heparinase [3, 7] or heparan sulfate (Figure S1h,i) cause dorsalization and alter *spry4* target gene expression. Taken together, these results indicate that specific extracellular HSPGs, but not cell-surface bound HSPGs, may directly regulate the range of Fgf8 signaling, potentially serving differing roles. Interestingly, for Dpp morphogen in the fly wing, distinct roles were recently observed in long range spreading and in local signaling at source cells, respectively [19].

Since only extracellularly localized HSPGs influenced Fgf8 signaling, we probed for a direct molecular interaction of Fgf8 with these molecules in the ECM in living embryos. Dual-color fluorescence cross-correlation spectroscopy (FCCS) involves monitoring fluorescence fluctuations of diffusing molecules from two spectrally different channels in a static Gaussian focal volume over time [36, 37]. Statistical auto-correlation of the signal in individual channels and cross-correlation in two channels can then be used to obtain absolute molecular concentration and quantify co-diffusion behavior (see Methods). We tested eGFP-tagged Fgf8 (Fgf8-eGFP) for cross-correlation with mRFP-tagged extracellular HSPGs (Figure 1a,b). All constructs (except Gpc2^ex^mRFP and Gpc5c^ex^mRFP) were found uniformly distributed in the ECM (Figure 1 c-f, Figure S2e-q). Exemplary FCCS curves are shown for controls and Gpc4^ex^mRFP *versus* Fgf8-eGFP pair, indicating the auto- and cross-correlation amplitude (Figure 1 g-i).

**Figure 1:**
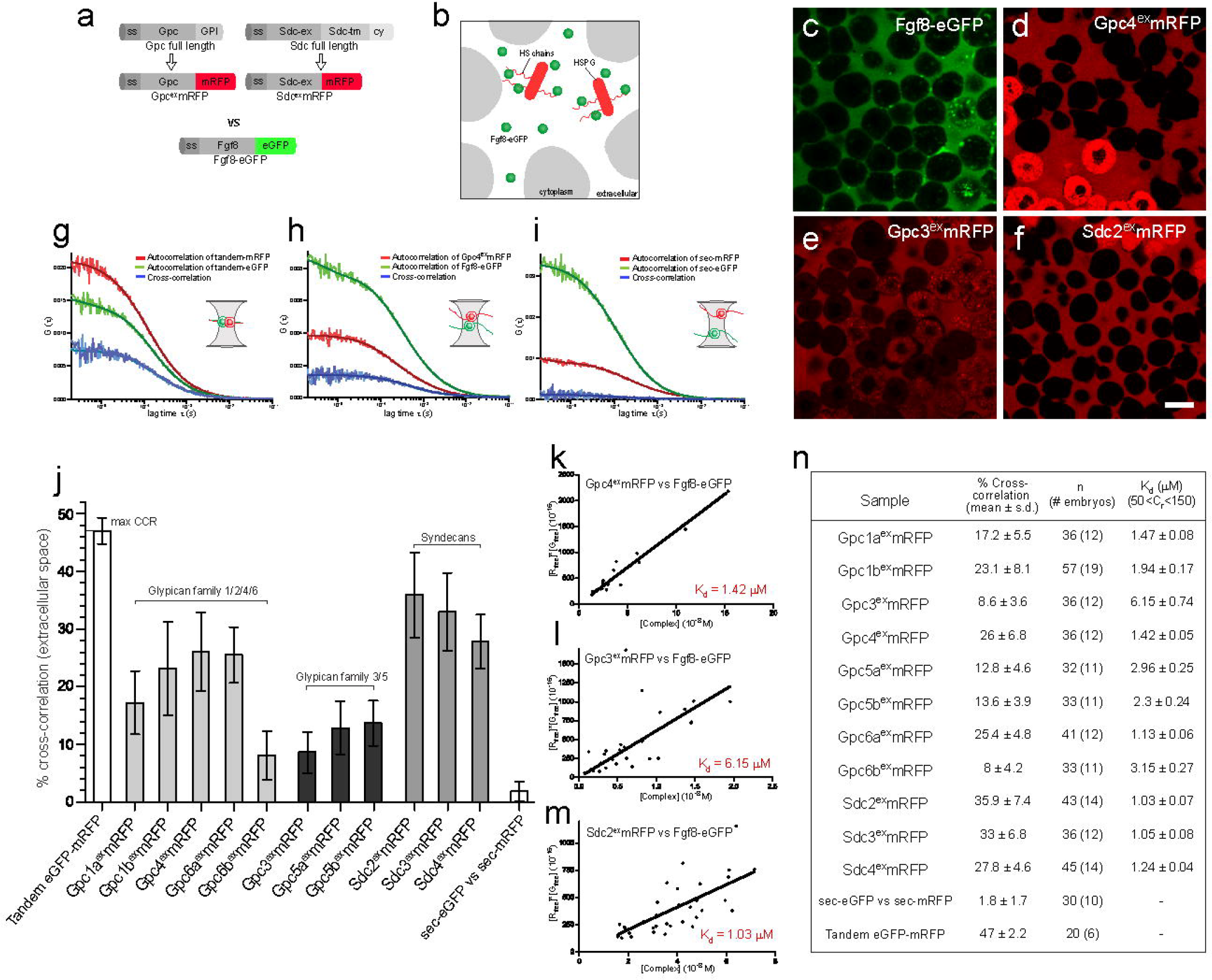
Fgf8 directly binds to glypican and syndecan HSPG in the extracellular space. (a) Illustration of constructs used for FCCS. GPI-anchor from Gpc and cytoplasmic and transmembrane domains from Sdc were replaced with mRFP, and subsequently used for FCCS with Fgf8-eGFP. ss: signal sequence, Gpc: glypican domain, GPI-GPI-anchor, Sdc-ex: syndecan extracellular domain, Sdc-tm: transmembrane domain, cy: cytoplasmic domain. (b) Schematic showing the site of measurement in the ECM and the potential interaction between Fgf8-eGFP (green) with extracellular HSPG (red). (c-f) Fusion proteins localize to the ECM in embryos. One member from each HSPG family is represented here: (c) Fgf8-eGFP, (d) Gpc4^ex^mRFP (Gpc family 1/2/4/6), (e) Gpc3^ex^mRFP (Gpc family 3/5) and (f) Sdc2^ex^mRFP. mRNA was injected at 32-cell stage and embryos imaged from the animal pole after 5 hpf. Scale bar: 20 μm. (g-i) Representative FCCS curves and schematic of molecular interactions in the Gaussian volume. Red and green curves represent the autocorrelation amplitude in red and green channel, respectively. Blue curves represent cross-correlation amplitude. (j) FCCS curve obtained for maximum cross-correlation (max. CCR) with tandem-eGFP-mRFP; schematic depicts strong association of fluorescent tags. (k) FCCS curve for Gpc4^ex^mRFP *versus* Fgf8-eGFP interaction; schematic represents partial binding. (l) Absence of cross-correlation between sec-eGFP and sec-mRFP, schematic represents no molecular interaction. (j) Percentage of cross-correlation for Fgf8-eGFP and various HSPGs and controls. The different HSPG families are indicated. A higher percentage of cross-correlation was measured for Sdc, followed by Gpc family 1/2/4/6 followed by family 3/5. Bar graph represents mean with s.d. The significance of the data was inferred using one-way ANOVA. All data points were highly significant compared to sec-eGFP vs sec-mRFP control, with p-value <0.0001. (k-m) Exemplary scatter plots for evaluation of effective dissociation constant (K_d_) for binding between Fgf8-eGFP and representative HSPGs: (k) Gpc4^ex^mRFP, (l) Gpc3^ex^mRFP and (m) Sdc2^ex^mRFP. K_d_ values are indicated in red. (n) Table showing FCCS measurements and corresponding K_d_ values for interaction of Fgf8-eGFP with different HSPGs in the ECM.

Percentage of cross-correlation (%CCR) is directly related to binding affinity and was obtained for Fgf8-eGFP and various extracellular HSPGs (Figure 1k-n). Embryos injected with mRNA encoding tandem-eGFP-mRFP as a positive control, showed a high %CCR (47 ± 2.2 % (mean ± s.d, n= 20, 6 embryos)), and very low %CCR was seen for sec-eGFP *versus* sec-mRFP as a negative control (1.8 ± 1.7 % (mean ± s.d, n= 30, 10 embryos)). When tested for cross-correlation with Fgf8-eGFP, the three Sdc: Sdc2^ex^mRFP, Sdc3^ex^mRFP and Sdc4^ex^mRFP showed higher cross-correlation with Fgf8-eGFP than Gpc proteins (Figure 1j,n). Among the Gpc’s, Gpc4^ex^mRFP had the highest %CCR, followed by Gpc6a^ex^mRFP and Gpc1a^ex^mRFP. This result complements our previous observation that extracellular Sdc, along with Gpc1a, Gpc4 and Gpc6a, influence Fgf downstream signaling (Figure S1e-i). Together, these data suggests that direct binding to specific extracellular HSPGs results in extended Fgf downstream signaling range.

Cross-correlation analysis was further used to evaluate *in vivo* dissociation constants K_d_ between Fgf8-eGFP and HSPGs (see Methods). Since there are multiple HSPGs interacting with Fgf8 in the extracellular space, the K_d_ obtained is an “effective” K_d_, reflecting more accurately the physiologically relevant values. K_d_ for Sdc and Fgf8 was between 1.03-1.24 μM, indicating strongest affinity amongst HSPGs (Figure 1k-m, Figure S3). Gpc3^ex^mRFP was the weakest binding HSPG with low %CCR and high K_d_ (%CCR= 8.6 ± 3.6 (mean ± s.d.); K_d_= 6.15 ± 0.74 μM). Overall, we observed that Fgf8 binding to individual extracellular HSPGs is a weak interaction (K_d_ in micromolar range), presumably reflecting the redundant nature of these molecules and the need for reversible binding and unbinding during Fgf8 propagation through the tissue. Moreover, our results support the restricted diffusion model of morphogen transport according to which the local diffusivity of a morphogen remains high but global diffusivity is slowed down due to interaction with extracellular regulators and receptors [3, 7, 38].

The affinity of Fgf8-eGFP for different HSPGs might depend on the core proteins, their HS chains or both. To test this, we generated HS-deficient HSPGs constructs (see Methods). The %CCR of Fgf8-eGFP was highly reduced for the tested HS-deficient HSPGs (Figure S4a,b). This implies that Fgf8-eGFP directly binds to HS chains in the ECM, which are likely to differ between different core-proteins. A previous genetic study showed that HS chains of HSPG are important for the retention of Fgf ligands around a cell’s perimeter [39]. Our observation extends this notion to include a requirement for HSPG core proteins. The binding affinity of Fgf8-eGFP for different HSPGs follows the order: Sdc >Gpc family 1/2/4/6 > Gpc family 3/5 (Fig 1n). Interestingly, the number of HS side chains is maximum in Sdc (3-6 SG repeats), followed by Gpc 1/2/4/6 (3-4 SG repeats) and Gpc 3/5 (1-2 SG repeats) [27, 40]. Hence an increasing number of HS chains, or a difference in HS-fine structure, may correlate with the differential affinity between Fgf8 and HSPGs.

We next tested the influence of HSPG binding on the diffusion of Fgf8-eGFP using autocorrelation-FCS [3]. In wild-type embryos injected with Fgf8-eGFP mRNA, fluorescence signal was localized in the ECM and receptors on the membrane (Figure 2a). Ectopic expression of Gpc4^ex^mRFP and Sdc2^ex^mRFP affected Fgf8-eGFP localization such that the fluorescence was stronger in the ECM and cell membrane binding could not be discerned (Fig 2b,c). FCS autocorrelation revealed longer diffusion times for Fgf8-eGFP upon binding (Figure 2f). Using a 2-component model for fitting the autocorrelation data (Methods), the diffusion coefficient (D) of Fgf8-eGFP fast component significantly reduced from D= 45 ± 9.8 μm^2^/s (mean ± s.d. (n= 64, 16 embryos)) in wild-type, to 33 ± 8 μm^2^/s (mean ± s.d. (n= 74, 18 embryos)) upon Gpc4^ex^mRFP injection, and 24 ± 3 μm^2^/s (mean ± s.d. (n= 29, 10 embryos)) upon Sdc2^ex^mRFP injection (Figure 2f,h). This reduction in diffusivity was not due to a higher extracellular viscosity because the D of sec-eGFP control did not change under similar conditions (Figure 2g,h). It was shown previously that a small fraction of Fgf8 molecules diffuse as slow component with D= 4 μm^2^/s [3, 7]. Upon over-expression of Gpc4^ex^mRFP and Sdc2^ex^mRFP, no change in the slow component was observed (Figure S4c). This indicates that the fast-diffusing Fgf8 is most likely the functionally relevant form of Fgf8 which is mobile and sensitive to the levels of extracellular HSPGs. It is likely that the slow component consists of Fgf8 bound to higher order structures in the ECM, which remains unaffected by transient HSPG expression.

**Figure 2:**
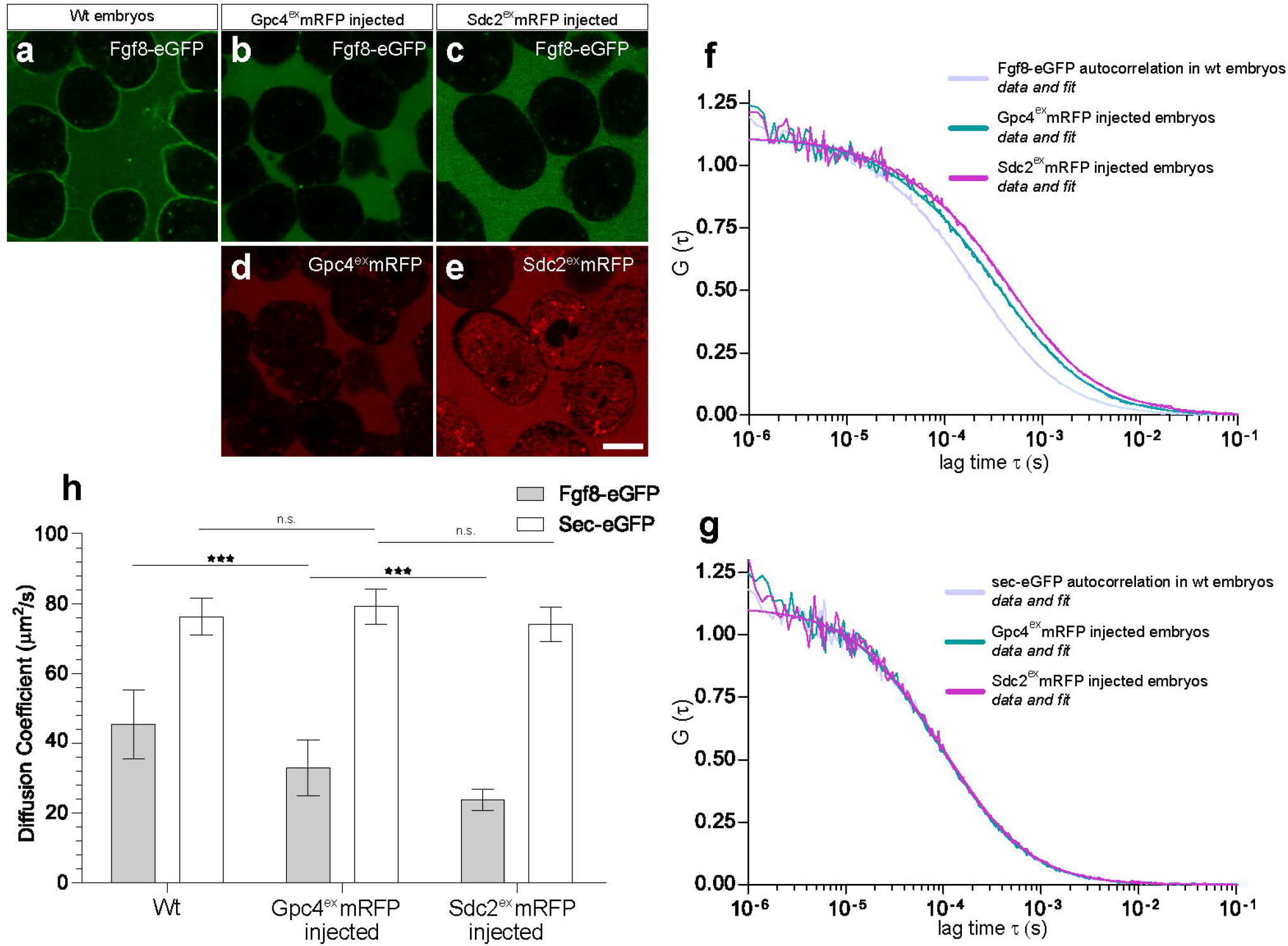
Binding to extracellular HSPG reduces Fgf8 diffusion. (a-c) Localization of Fgf8-eGFP in wild-type embryos (a) and embryos injected with Gpc4^ex^mRFP mRNA (b, d) and Sdc2^ex^mRFP mRNA (c, e). Note the extracellular retention of Fgf8-eGFP in b and c. Scale bar: 10 μm. (f) Autocorrelation curve of Fgf8-eGFP in wild-type and embryos over-expressing Gpc4^ex^mRFP and Sdc2^ex^mRFP. The curve shifts towards longer diffusion times upon binding to Gpc4^ex^mRFP and Sdc2^ex^mRFP, indicating a reduction in mobility. (g) Autocorrelation of sec-eGFP is not affected by the presence of excess Gpc4^ex^mRFP and Sdc2^ex^mRFP in embryos. The autocorrelation function was fit to a 2-component model for Fgf8-eGFP and 1-component model for sec-eGFP. (h) Diffusion coefficient of Fgf8-eGFP (grey bars) and sec-eGFP (white bars) in embryos over-expressing Gpc4^ex^mRFP and Sdc2^ex^mRFP. Diffusion coefficient of Fgf8-eGFP is reduced, but sec-eGFP does not change. Bar graph represents mean with s.d. Statistical significance was inferred using one-way ANOVA.

Enhanced Fgf downstream signaling upon extracellular binding could be due to HSPG-mediated further spreading of Fgf8, or due to a greater stability of Fgf8 in the ECM and hence longer persistence. To test this, we visualized the Fgf8 gradient in early embryos (4 hpf) in the presence and absence of extracellular binding. Following a previously established protocol [3], fluorescence intensity of Fgf8-eGFP was quantified at different ECM positions away from a local source of production (Figure 3a). The concentration profile is described by the decay length (λ) (see Methods); for wild-type embryos, we obtained λ= 202 ± 7.1 μm (mean ± s.d, n= 25 embryos) (Figure 3b). Ectopic expression of Gpc4^ex^mRFP and Sdc2^ex^mRFP significantly reduced Fgf8-eGFP decay length to 178 ± 6.7 μm (mean ± s.d, n= 18 embryos) and 179 ± 8.2 μm (mean ± s.d, n= 17 embryos), respectively (Figure 3b,c). This result indicates that direct extracellular binding, which reduces Fgf8-eGFP diffusion, leads to a steeper Fgf8 gradient and enhanced downstream signaling during gastrulation could, for instance, be a result of a greater protein stability. Using a morpholino-based knock-down approach, we also tested the effect of Gpc4 loss-of-function on Fgf8-eGFP gradient shape. Injection of translational and splice-junction morpholinos against Gpc4 (Gpc4-MO), recapitulated the *gpc4* (*knypek*) mutant phenotype [41] and led to a reduction in the fluorescent signal of Gpc4^ex^mRFP injected embryos (Figure S4d,e), indicating a specific reduction in *gpc4* translation. When examined for Fgf8-eGFP gradient profile, Gpc4 knock-down resulted in an increased decay length and a shallower gradient (λ= 270 ± 15 μm (mean ± s.d, n= 20 embryos)) (Figure 3d,f). The gradient profile was unaltered after injection of control morpholino, λ= 210 ± 9.8 μm (mean ± s.d, n= 14 embryos) (Figure 3e). Together these results suggest that HSPGs, and specifically Gpc4, regulate Fgf8-eGFP gradient shape by restricting its diffusion in the ECM. Moreover, since we observe that altering ECM composition affects the gradient outcome, there needs to be a proper balance between freely diffusing *versus* ECM bound Fgf8 molecules to obtain the optimum gradient length.

**Figure 3:**
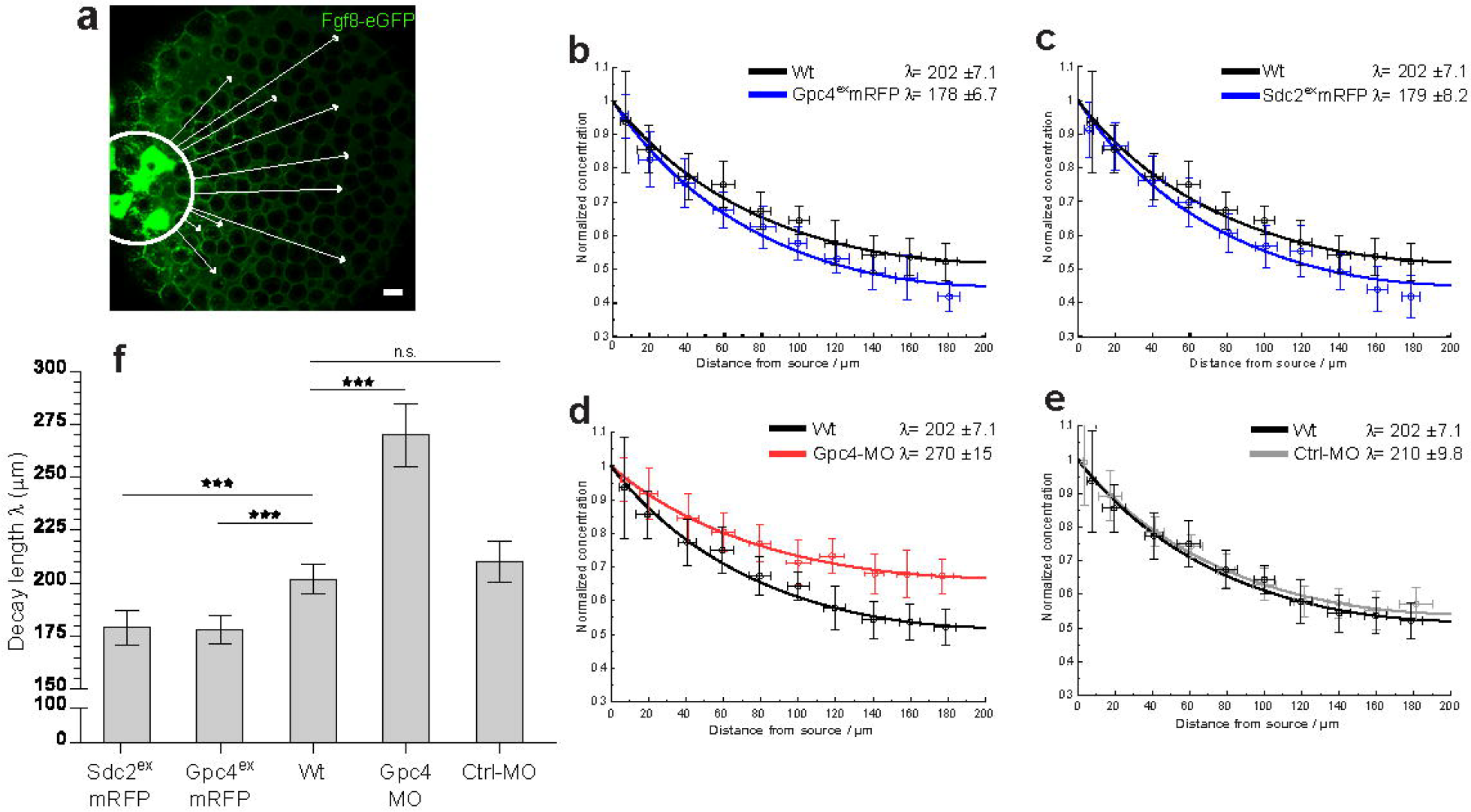
Fgf8 gradient is influenced by changing the composition of extracellular matrix. (a) Confocal image showing extracellular Fgf8-eGFP gradient from a restricted source of cells (white circular arc). The source was created by injecting Fgf8-eGFP mRNA in 1-blastomere at the 32-cell stage and confocal images acquired after 4 hours. The fluorescent intensity was measured at different positions (white arrows) away from this source in the free extracellular space. Scale bar: 20 μm. (b-e) Normalized concentration gradient binned in 20 μm intervals and fit to a radial symmetry model to obtain the decay length *λ* [3]. Comparison of Fgf8-eGFP gradient profile in wild-type embryos (black) and embryos expressing Gpc4^ex^mRFP (b, blue) and Sdc2^ex^mRFP (c, blue). The gradient becomes steeper due to extracellular binding and reduction of Fgf8-eGFP diffusion. (d) The gradient assumes a shallower profile after morpholino (MO) mediated knockdown of Gpc4 (red). (e) Injection of a control morpholino does not influence Fgf8-eGFP distribution (grey, n= 14). (f) Comparison of decay length (*λ*) under specified conditions of ECM modification. Graph represents mean with s.d. One-way ANOVA was used to evaluate statistical significance.

Since HS chains are essential for Fgf8-Fgf receptor (FgfR) membrane interaction [21, 23], we tested whether extracellular HSPGs regulate and/or interfere with this interaction. Dual color membrane scanning FCS was used to quantify interactions at the cell membrane, which is better suited for slowly moving molecules on the membrane (Figure 4a-c, Methods) [16, 42]. We obtained high cross-correlation (%CCR= 34.4 % ± 2.9 (mean ± s.e.m., n= 21)) for the interaction of Fgf8-eGFP with its receptor (FgfR1), as reported earlier (Figure 4d) [16]. We tested the affinity of Fgf8-eGFP for its receptor in the presence of excess untagged extracellular Gpc4 (Gpc4^ex^) and in its absence after Gpc4-MO injection (Figure 4f). Notably, the affinity of Fgf8-eGFP for its receptor did not change upon binding to extracellular Gpc4^ex^, compared to the control (Figure 4g). Additionally, in Gpc4-MO injected embryos this membrane interaction was also not affected (Figure 4g). This result suggests that the affinity of Fgf8-eGFP for its receptors is independent of extracellular HSPGs, although a possible contribution of other HSPG molecules remains to be tested.

**Figure 4:**
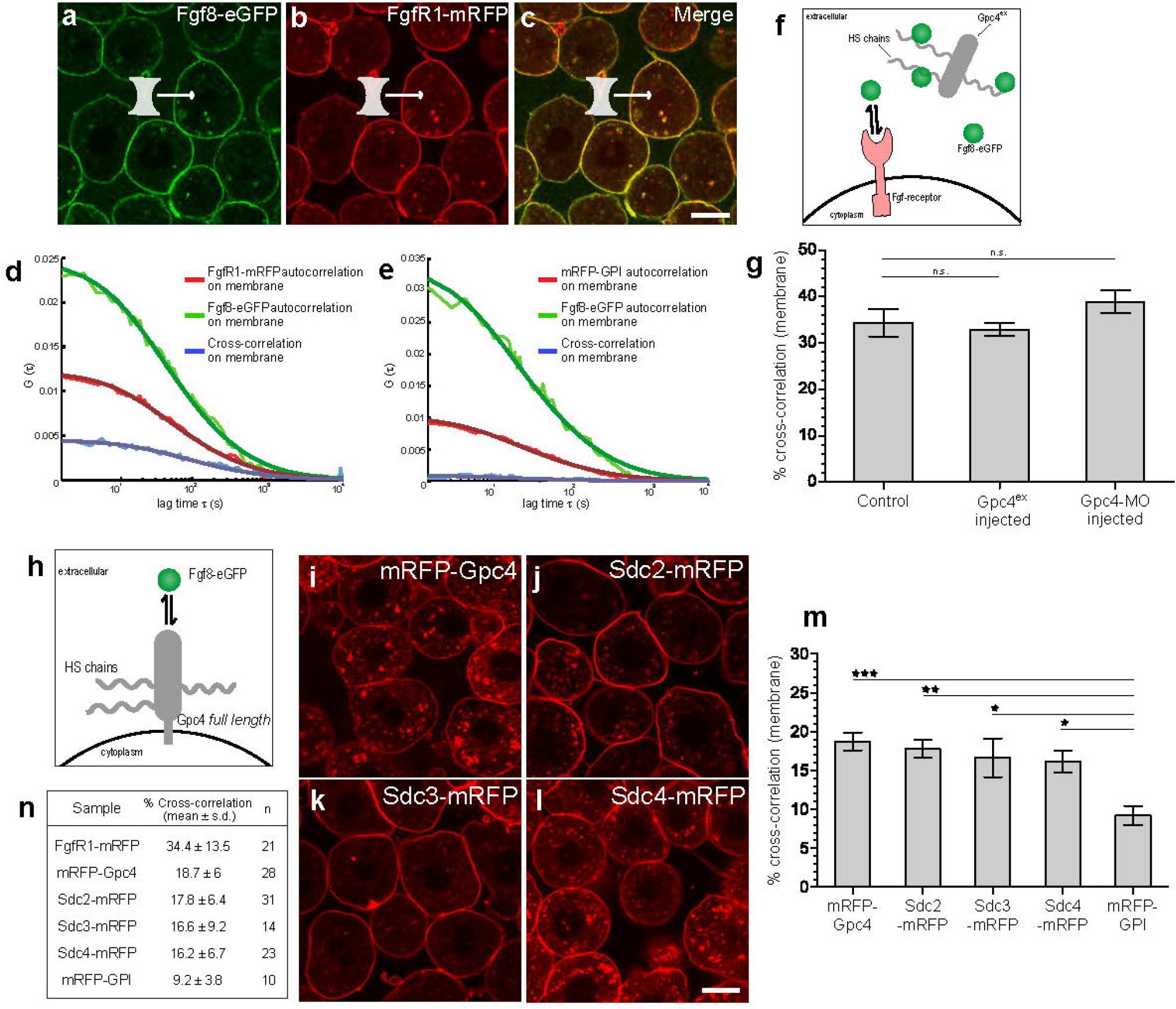
Fgf8 and Fgf-receptor interaction is regulated by cell membrane bound and not extracellular HSPG. In dual color scanning FCS fluorescent intensity fluctuations of membrane localized molecules are monitored and used for cross-correlation analysis. The Gaussian focal volume is scanned across the membrane to obtain photon counts. (a-c) Membrane localization of Fgf8-eGFP (a), FgfR1-mRFP (b) and merged image (c) are shown for illustration. Scale bar: 10 μm. (d-e) Exemplary cross-correlation curves obtained after scanning FCS on the membrane. (d) High amplitude of cross-correlation (blue) was observed for FgfR1-mRFP and Fgf8-eGFP binding. (e) Cross-correlation was negligible for mRFP-GPI control and Fgf8-eGFP interaction on the membrane. (f) Schematic showing aim of the experiment to directly monitor Fgf8-eGFP and FgfR1-mRFP interaction after addition or removal of Gpc4 in the ECM. (g) Fgf8-eGFP affinity for FgfR1-mRFP did not change significantly upon addition of excess Gpc4^ex^ (n= 21) or morpholino-mediated knockdown of Gpc4 (n= 26), as compared to wild-type control (n= 21). Graph represents mean with s.e.m. One-way ANOVA was performed on the dataset to infer statistical significance. (h) Illustration depicting aim of the experiment to measure direct binding between Fgf8-eGFP and cell-surface attached HSPGs. (i-l) Membrane localization of different HSPG constructs. mRFP was attached at the N-terminal for Gpc4 (i) and C-terminal for Sdc2 (j), Sdc3 (k) and Sdc4 (l). Scale bar: 10 μm. (m) Scanning FCS revealed that Fgf8-eGFP directly binds cell-surface attached mRFP-Gpc4, Sdc2-mRFP, Sdc3-mRFP and Sdc4-mRFP with similar affinities. Graph represents mean with s.e.m. One-way ANOVA and Dunnett’s multiple comparison tests were performed to test for significance. (n) Cross-correlation values obtained from membrane scanning FCS between Fgf8-eGFP and the indicated constructs.

Although we did not observe extracellular HSPGs to interfere with cell-surface binding, we argued that cell-membrane HSPGs might provide the necessary HS chains for Fgf8-FgfR ternary complex [43] formation. To investigate this, we measured the interaction of Fgf8-eGFP with cell-surface attached Gpc4 and Sdc proteins (Figure 4h). Full-length membrane attached mRFP-Gpc4 and Sdc (Sdc2-mRFP, Sdc3-mRFP, Sdc4-mRFP) showed uniform membrane localization (Figure 4i-l). Membrane scanning FCS revealed a similar binding affinity for all tested HSPGs, such that %CCR for mRFP-Gpc4 was 18.7 ± 6.3 (mean ± s.e.m., n= 28), for Sdc2-mRFP: 17.8 ± 6.4 (mean ± s.e.m., n= 31), for Sdc3-mRFP: 16.6 ± 9.2 (mean ± s.e.m., n= 14) and Sdc4-mRFP: 16.2 ± 6.7 (mean ± s.e.m., n= 23) (Figure 4m,n). mRFP-GPI was used as a negative control with %CCR= 9.2 % ± 3.8 (mean ± s.e.m., n= 10) (Figure 4e). It is interesting to note that positive binding was seen only at higher concentrations of cell-surface bound HSPGs (100-200 pg mRNA), whereas Fgf8-eGFP and FgfR1-mRFP interaction could be detected already at 30 pg of FgfR1-mRFP mRNA. Therefore, the binding interaction between Fgf8 to HSPGs on the membrane appears weaker than its receptor interaction. Nevertheless, our results support the hypothesis that HS chains for Fgf-FgfR interaction are presented by cell-surface bound HSPGs. Interestingly, when we analyzed the Fgf8-eGFP extracellular gradient upon over-expression of cell-membrane bound Gpc4, the decay length was reduced similarly to over-expression of Gpc4^ex^mRFP in the ECM (Figure S4f,g). This implies that cell membrane attached Gpc4 does not directly influence extracellular Fgf8 distribution, but does so only after secretion from the cell surface. For instance, cell-surface bound HSPGs might help prevent FgfR-mediated endocytosis and degradation, as recently suggested for Dpp [44]. Overall, our *in vivo* cross-correlation results suggest a dual-role of HSPGs: (i) retention of Fgf8 ligand in the ECM, mediated by extracellular HSPGs and (ii) cell-surface co-receptor function facilitating the formation of ternary receptor-ligand complex, mediated by cell-surface attached HSPGs [21-23, 43].

To complement our observations, we used correlative light and electron microscopy to directly visualize the distribution of Fgf8-eGFP around cell membranes (Figure 5, Methods). Interestingly, gold-labeled Fgf8-eGFP molecules were observed at different positions with respect to the membrane: >500 nm away from membrane (Figure 5e), 10-50 nm away and directly on the membrane (Figure 5f, Figure S5). These species most likely correspond to freely diffusing, ECM-bound and receptor-bound Fgf8-eGFP molecules, respectively. The ECM-bound ‘immobile’ fraction of Fgf8 has not been visualized previously and confirms our hypothesis that ECM acts as a reservoir for the local retention of Fgf8 ligand.

**Figure 5:**
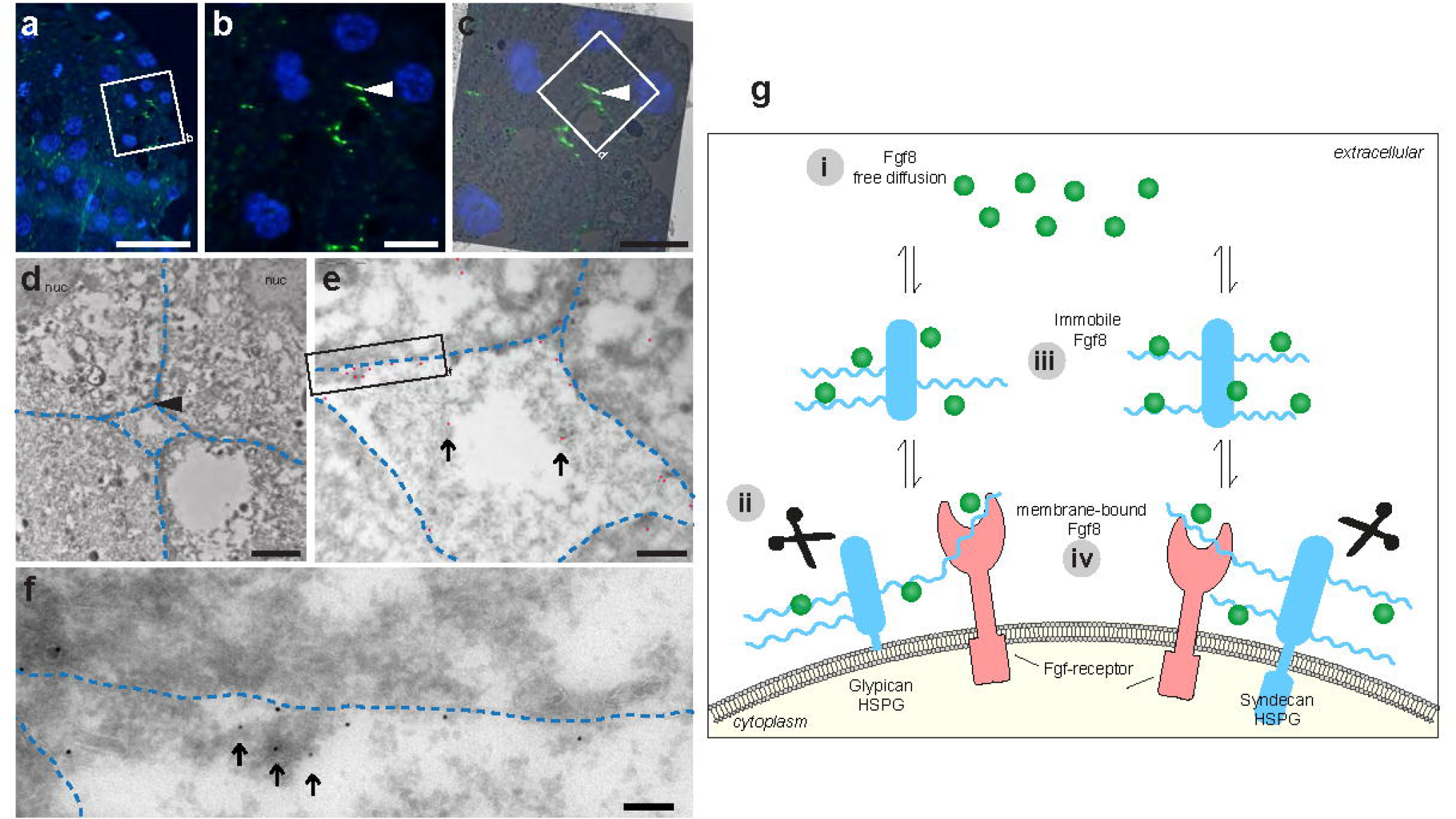
Electron microscopy-based visualization of Fgf8 around the cell membrane. (a-f) Correlative light electron microscopy to study localization of Fgf8-eGFP at high resolution around cell membranes. (a-c) Light microscopy image of a 70 nm cryo-section, immunostained with rabbit anti-eGFP protein A gold, goat anti-rabbit Alexa488 and DAPI (nucleus). From fluorescent images, areas showing strong GFP labeling (filled arrowhead) and intact membrane/extracellular space were imaged further in the TEM. (d-f) Extracellular space enclosed by cell membranes on four sides, revealed localization of 10 nm gold-labeled Fgf8-eGFP molecules. (e) Arrows point at freely diffusing Fgf8-eGFP particles located ∼1 μm away from cell membranes. (f) A close-up of the membrane (highlighted in blue) shows that even in the vicinity of the membrane, particles are distributed at a distance of 10-50 nm away from the membrane (arrows), representing both membrane and ECM bound molecules. Scale bar (a): 50 μm, (b): 10 μm, (c) 10 μm, (d): 2 μm, (e): 500 nm, (f): 100 nm. (g) Proposed model for the regulation of extracellular Fgf8 gradient by HSPG. (i) Most Fgf8 molecules (green) traverse the embryo by free diffusion. (ii) Glypicans and syndecan heparan sulfate proteoglycans (blue) are constitutively shed from the cell surface into the ECM. (ii) Due to binding with extracellular HSPGs, Fgf8 gets trapped near the cell surface. HSPGs bind Fgf8 mainly via their HS side chains. This transient binding forms a relatively immobile Fgf8 fraction and is essential to obtain the optimal gradient length. (iv) Formation of a ternary complex with Fgf-receptor (red), Fgf8 and HS sugar chains, involves cell-surface bound HSPGs. The extracellular secreted HSPGs do not contribute to cell-membrane interactions.

In this study we show, by using single molecule FCS in living embryos at the time of ongoing patterning and signaling decisions, that in addition to free diffusion, Fgf8 distribution is directly regulated by binding to HSPGs in the ECM. Fgf8 binds with differential low-range affinity to Gpc and Sdc HSPGs in the ECM via their HS chains (Figure 1). We observed that the extracellular concentration of these molecules directly regulates Fgf8 diffusion and gradient shape (Figure 2,3) and that membrane-bound and extracellular matrix pools of HSPGs serve different roles (Figure 4). Correlative light and electron microscopy further revealed that Fgf8 is localized on the membrane as well as several nanometers away from the membrane (Figure 5a-f). Together with evidence of direct molecular binding, this observation indicates that HSPGs can trap Fgf8 near the cell surface. Our results also strongly support the existence of an immobile fraction of Fgf8, in complex with HSPGs, which acts as an intermediate between freely diffusing and membrane-bound molecules (Figure 5g). During gastrulation, a stage dominated not only by ongoing Fgf8 signaling and patterning, but also massive cell movements and rearrangements, the immobile fraction might play an important role in regulating the Fgf8 concentration that is experienced by cells. Thus, we propose that the distribution of Fgf8 is controlled at two levels: through a steady-state gradient in the animal-vegetal axis of the embryo whose formation is dominated by fast diffusion; and a concentration effect around the cells perimeters which is likely to be in equilibrium with receptor-bound Fgf8.

## Author Contributions

The project was designed and supervised by MB, in discussion with MG. Zebrafish experiments, including FCS experiments and data analysis, were done by MG; membrane scanning FCS experiments and interpretation by FH, MG and PS; embryo injection experiments, in situ hybridisation by MG, SK and FK; and electron microscopy by TK, using samples provided by MG. MG and MB wrote and edited the manuscript. All authors approved the manuscript.

## Acknowledgements

We are grateful to Andy Oates and Christian Bökel for excellent discussions concerning the project. We thank past and present members of the Brand lab for generous support and many helpful and intense discussions, S. Kretschmar for technical assistance, as well as M. Fischer and J. Michling for excellent zebrafish care, and the Light Microscopy Facility, a core facility of CMCB at Technische Universität Dresden, especially Markus Burkhardt and Wolfgang Staroske for help with analysis of FCS measurements. We also thank Stefan Hans, Gokul Kesavan and Markus Burkhardt for critical reviewing of the manuscript. Funding: We gratefully acknowledge the support through project grants to MB by the German Research Foundation (Deutsche Forschungsgemeinschaft, project numbers BR 1746/6-2, BR 1746/11-1, and BR 1746/3), a Cluster of Excellence ‘Physics of Life’ seed grant, and institutional funds from TU Dresden, to MB. Further support was by the DFG Transregio 67 (project 387653785), the DFG SPP 2084 μBone (project KN 1102/2-1) and the EXU transCampus funding program (tC2020_02_MED) to FK. The work at TU Dresden is co-financed with tax revenues based on the budget agreed by the Saxonian Landtag.

## Material and Methods

### Fish maintenance

Zebrafish (*Danio rerio*) were raised and maintained according to methods described previously [45]. Embryos were obtained by natural spawning and staged according to morphology [46]. For confocal imaging and FCS measurements, embryos at the sphere stage were mounted in 1% low melting-point agarose and maintained in methylene free E3 buffer [45] throughout measurements. All animal maintenance was carried out in accordance with animal welfare laws of the Federal Republic of Germany (Tierschutzgesetz) that were enforced and approved by the competent local authority (Landesdirektion Sachsen; protocol numbers TVV21/2018; DD24-5131/346/11 and DD24-5131/346/12).

### Generation of Plasmid Constructs

All constructs for mRNA injection were cloned in pCS2+ plasmid. The plasmids for the following constructs were obtained from the respective literature: Fgf8-eGFP and sec-eGFP [3], mRFP-Gpi and FgfR1-mRFP [16] and full length glypicans [27]. Syndecan-2 and syndecan-4 full length was kindly provided by J. R. Whiteford. Full length Sdc3 was amplified from zebrafish cDNA, using primers listed in Methods Table 2. Extracellular glypican and syndecan constructs (Gpc^ex^mRFP and Sdc^ex^mRFP) were generated by cloning the glypican domain and syndecan ectodomain in front of mRFP, usually spaced by a serine-glycine linker. Primers are listed in Methods Table 1. For tandem-eGFP-mRFP, eGFP was inserted in front of mRFP in pCS2+ vector. For generation of full-length mRFP-Gpc4, mRFP was placed in between the signal sequence and glypican domain. Full length mRFP-tagged syndecans were created by tagging mRFP to the cytoplasmic C-terminus (Methods Table 2).

**Methods Table 1:**
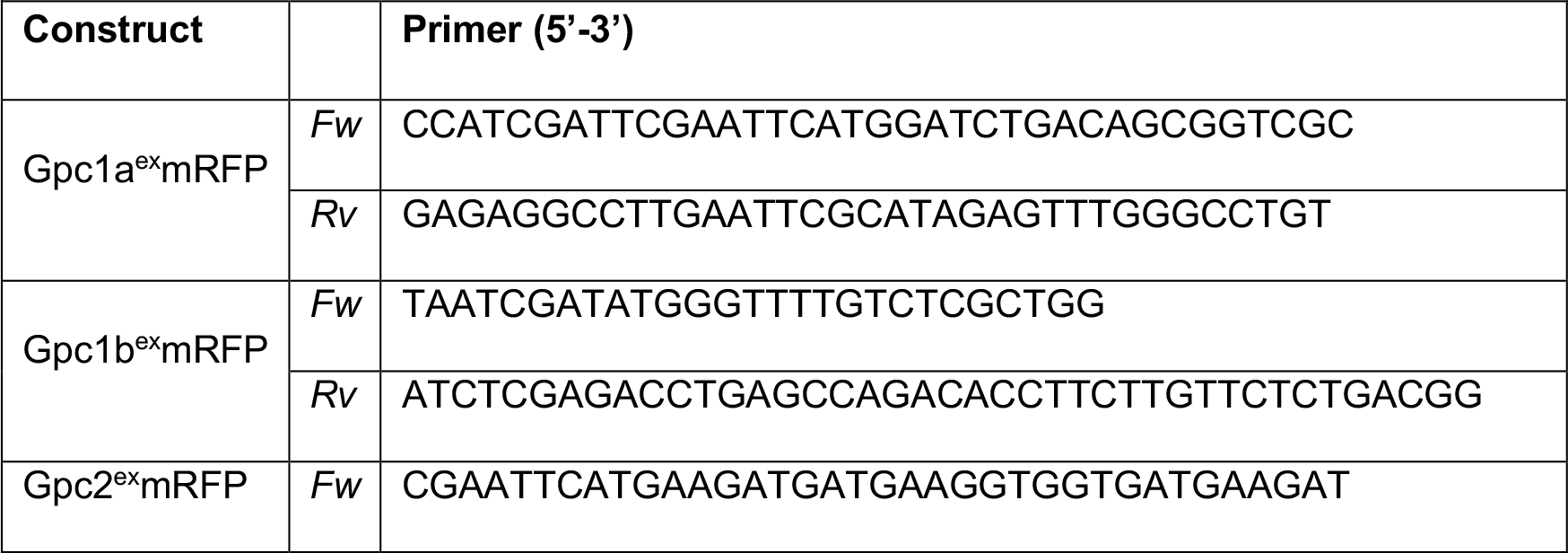
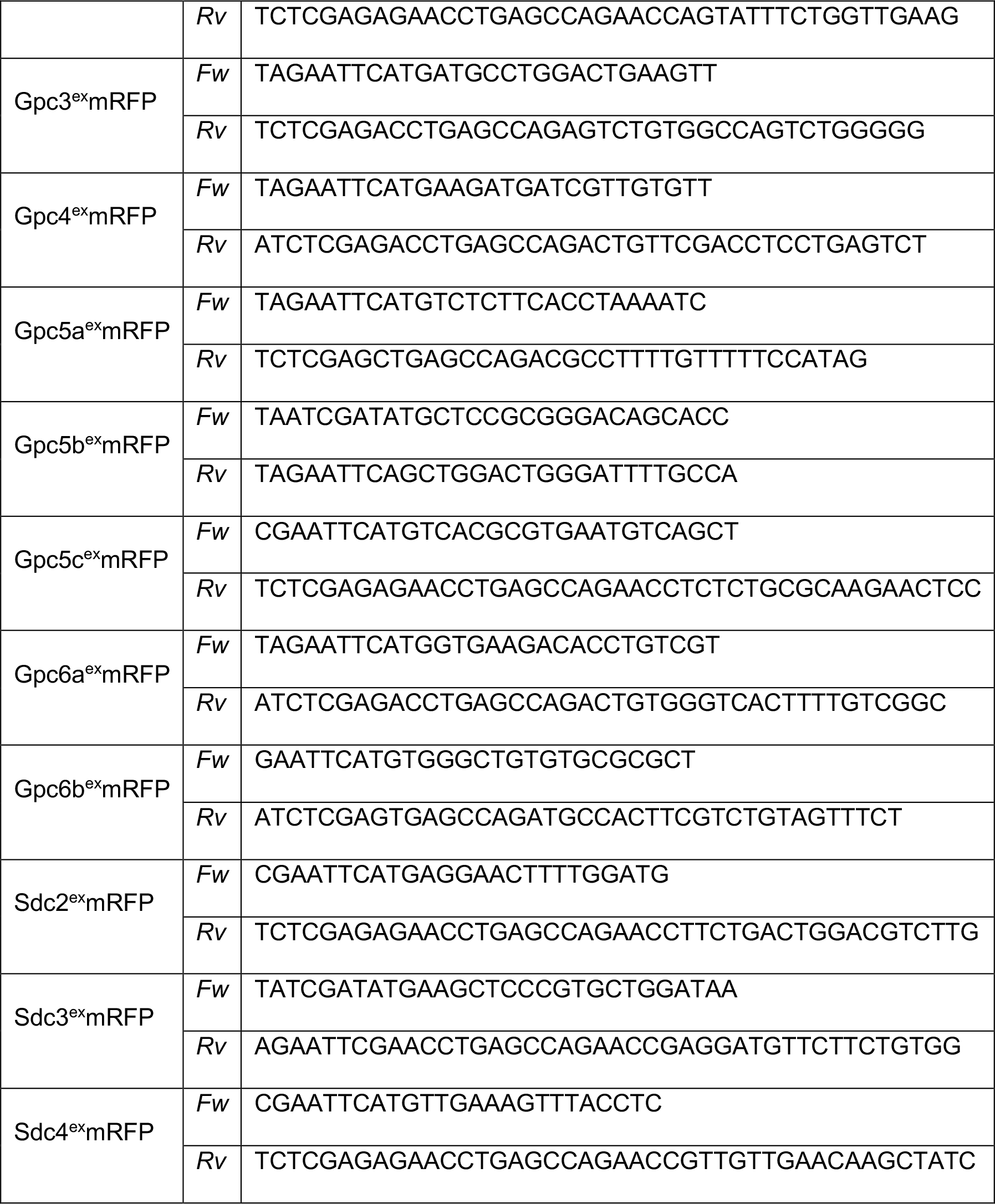
Primers used for cloning extracellular mRFP-tagged HSPG.

**Methods Table 2:**
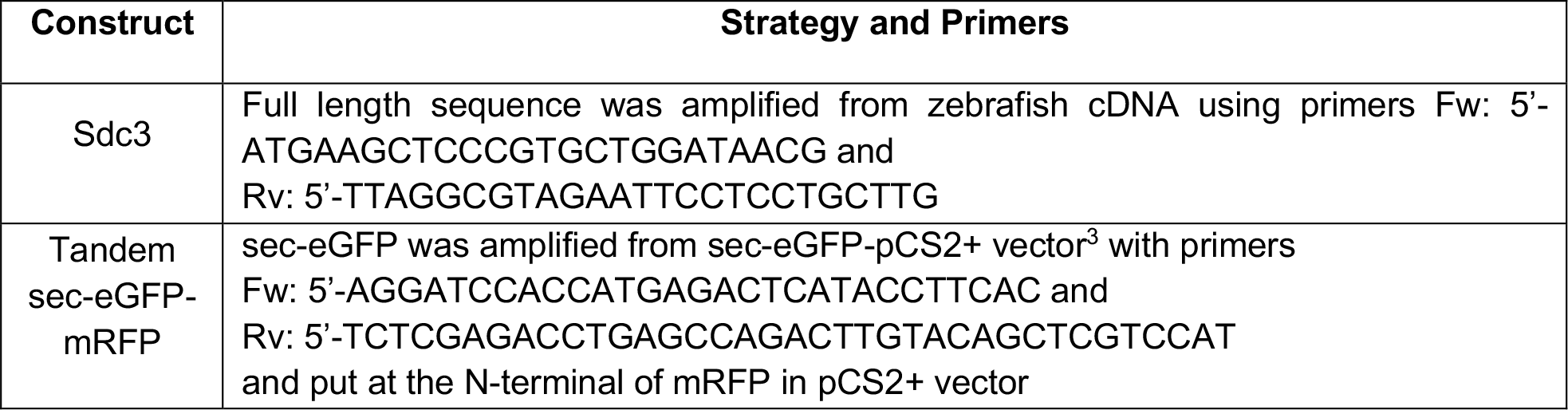
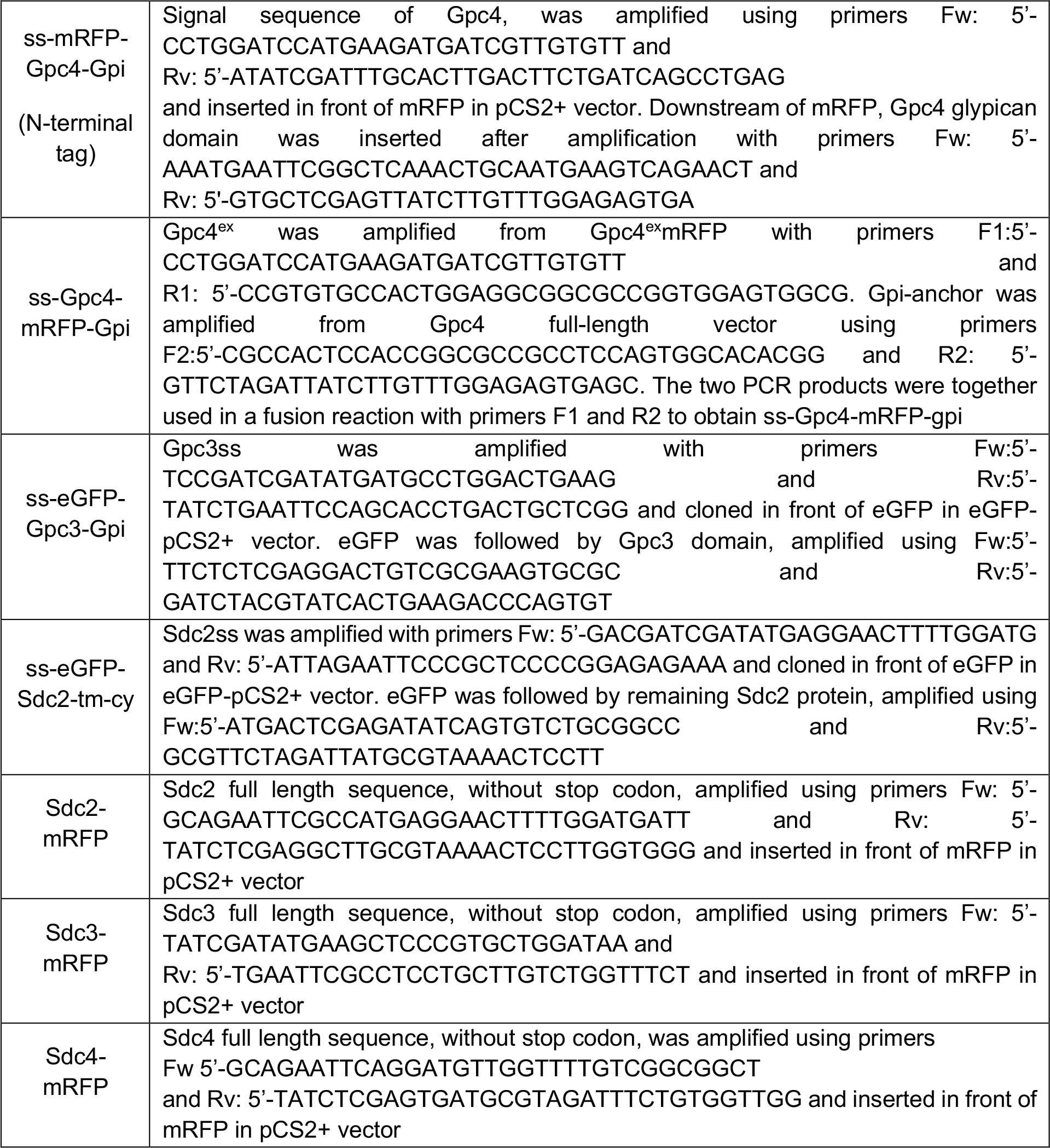

Site-directed mutagenesis: HS-deficient HSPG constructs were generated by mutating serines to alanines in serine-glycine repeats of the core proteins [47]. PCR mediated site-directed mutagenesis was used for multiple rounds to achieve this. Forward (*M*_*f*_) and reverse (*M*_*r*_) mutagenic primers were designed containing the mutation and 12 flanking complementary bases on both sides. The mutagenesis was carried out in three PCR steps: 1) Gene specific forward (*G*_*f*_) and mutagenic reverse primer; 2) Mutagenic forward and gene specific reverse (*G*_*r*_) primer; 3) Fusion reaction using gene specific forward and reverse primers and a mixture of the above two amplicons as template. For syndecans, mutagenesis was carried out in several subsequent mutagenesis PCR reactions. Methods Table 3 lists all primers used.

**Methods Table 3:**
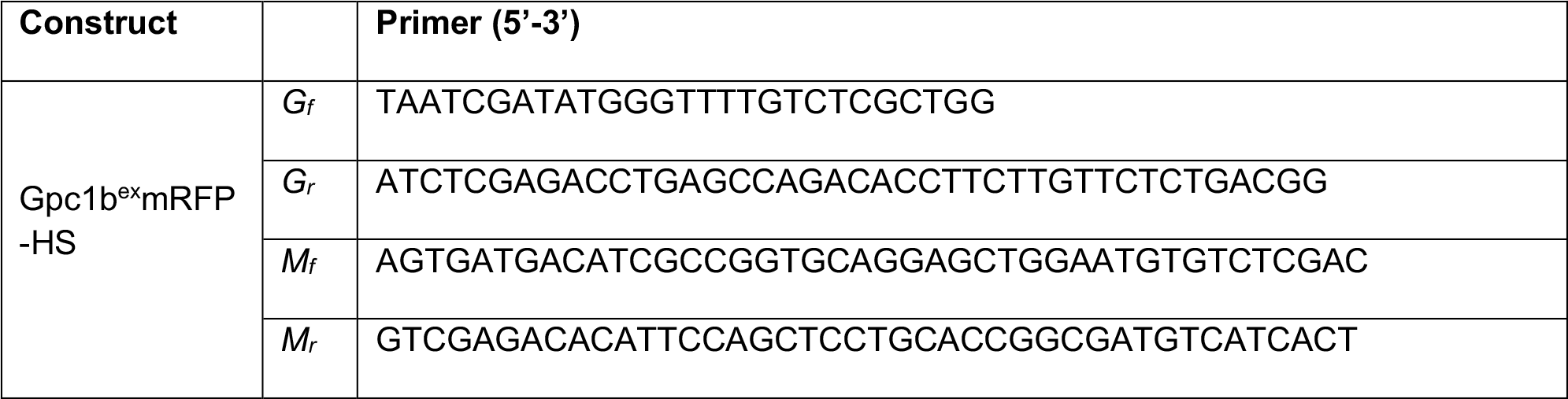
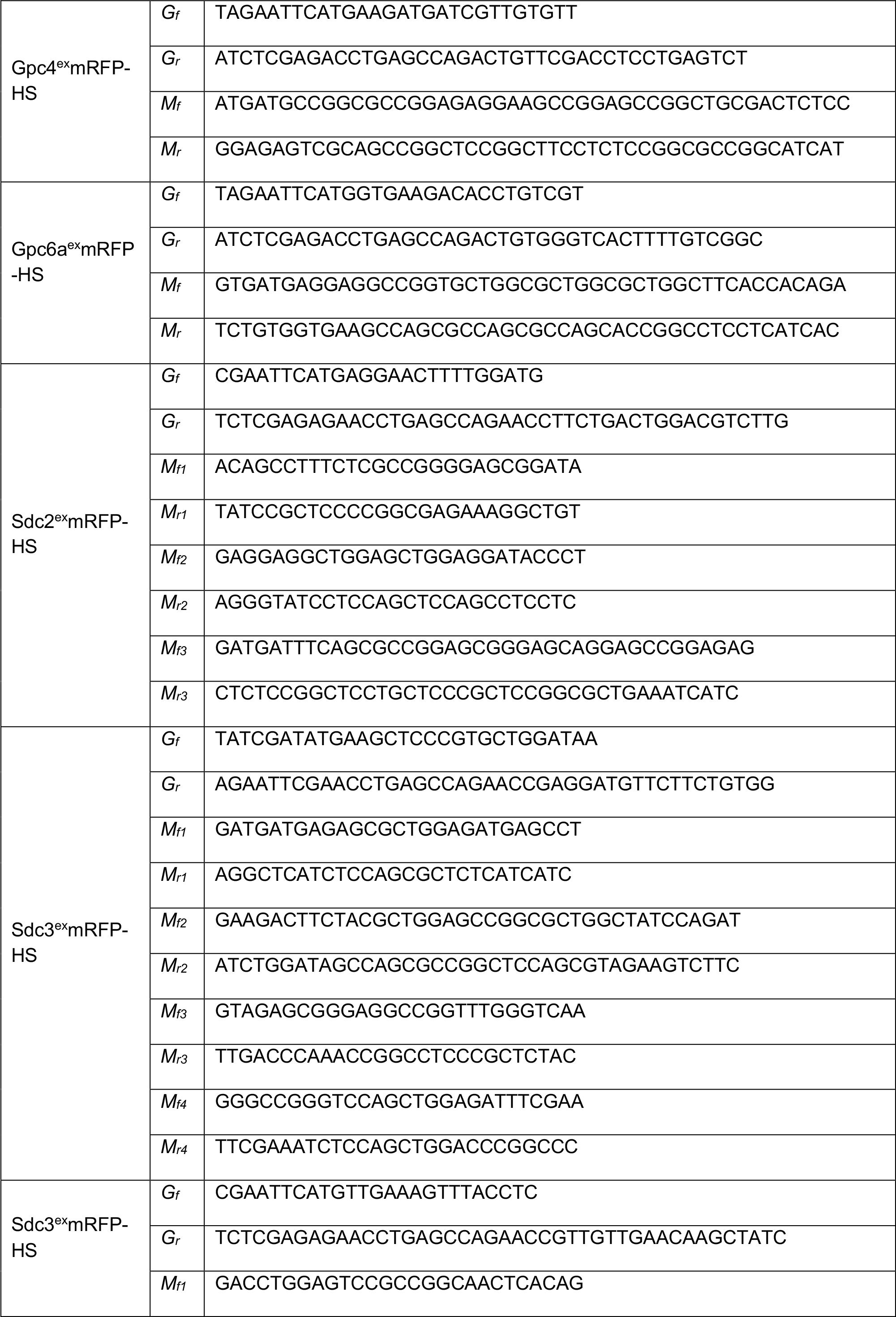
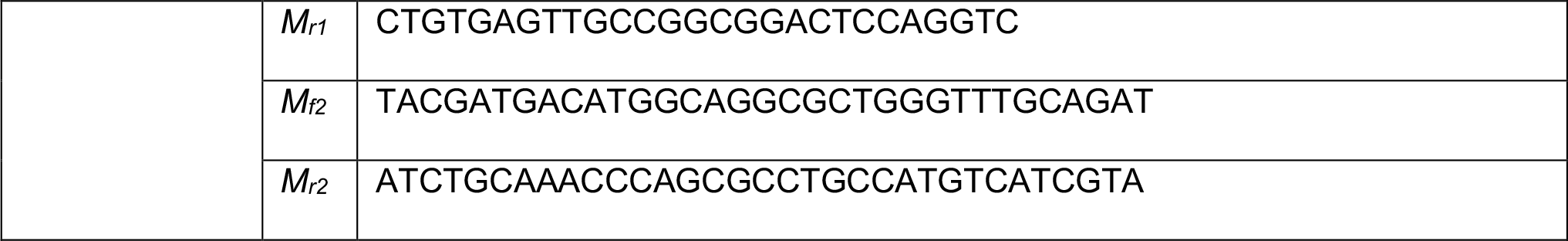
Primers used for mutating HS attachment sites in HSPG.

### Molecular Methods

mRNA for injection was prepared using mMESSAGE mMACHINE kit (Ambion). For over-expression of HSPG, 50-100 pg of required mRNA was injected in the cytoplasm of single-celled embryos. Fgf8-eGFP mRNA (20 pg) was injected in a single-blastomere at the 32-cell stage to generate a local source of production.

Morpholinos were obtained from Gene tools with the following sequence: Gpc4-MO1 (ATG block): 5’-CCACGACAGGTGTCTTCACCATCCC, Gpc4-MO2 (Splice block): ATAAAATCAGTGTAACTCACCCTGC.

For *in situ* hybridization, embryos were raised until 60 % epiboly stage and fixed in 4 % PFA. Anti-sense probe synthesis (*spry2, spry4, etv4* and *fgf8*) and *in situ* hybridization were carried out according to a previous protocol [10].

### FCS experimental set-up

Measurements were performed in the extracellular space in embryos between sphere- and 50% epiboly-stages, usually within 30 μm below the enveloping layer. Both auto- and cross-correlation measurements were performed with the Zeiss LSM780/Confocor 3 microscope and C-Apochromat 40x objective (N.A. 1.2). For auto-correlation of Fgf8-eGFP, the sample was excited with the 488 nm Argon laser line (laser power 8 μW), and the emitted photons were collected using an avalanche photodiode. The pinhole and objective correction collar were first calibrated using Alexa488 free dye in solution and the pinhole diameter set at 1 A.U., corresponding to 35 μm in the LSM780 microscope. Acquisition was done for 10*10 sec and an average was used for analysis after manually eliminating irregular curves. The data was analyzed using a 3-D free diffusion model of 2-components in Matlab (Mathworks) program (Jonas Ries [3]). Structural parameter for analysis was fixed at 6, eGFP triplet fraction (F_trip_) at 10 % and triplet blinking time (τ_trip_) at 30 μs, as determined from calibration of Alexa488 and eGFP in solution [3, 48].

For cross-correlation, samples were excited with the 488 nm and 561 nm Argon laser lines with laser powers 8 and 7.6 μW, respectively. Pinhole diameter was set at 1 A.U. for the longer wavelength (40 μm in LSM780 microscope). Calibration of the focal volume was done using 100 nM Alexa488 and 200 nM CF568 dyes in solution. The auto- and cross-correlation curves were fit using a 3-D free diffusion model in Zeiss Zen2012 software. The range of count rates obtained was 20-200 kHz in the green channel and 100-500 kHz in the red channel. Counts per molecule (cpm) were found to be stable on different days from 4-6 kHz for both channels.

For background correction, fluorescence intensity was recorded in both red and green channels in un-injected embryos. This value was subtracted from the count rate of all samples before cross-correlation analysis. For cross-talk correction, fluorescence signal was recorded in both channels after Fgf8-eGFP only injection. The cross-talk factor (β=*F*_*r*_/*F*_*g*_) was determined and the % cross-correlation curves was re-calculated including this factor.

### Fluorescence auto-correlation (FCS) analysis

Fgf8-eGFP auto-correlation analysis was performed as previously described [3]. The fluorescence auto-correlation function G(τ) is defined as the time-dependent decay in fluorescence fluctuation intensity and described as:

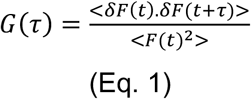

where < > denotes the time average and τ is the lag time. The fluorescent intensity fluctuations at time t (δF(t)) are compared to fluctuations at time (t+τ) and the signal is averaged over time. The concentration of fluorophores in the focal volume can be obtained from the amplitude of auto-correlation. At τ = 0, assuming the fluorescence intensity is proportional to the number of molecules N (which follow a Poisson distribution), then in eq.1,

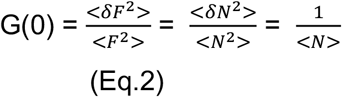

The measured auto-correlation curves were fit to the 3D free-diffusion model including eGFP triplet blinking. For a single diffusing species, this is given by:

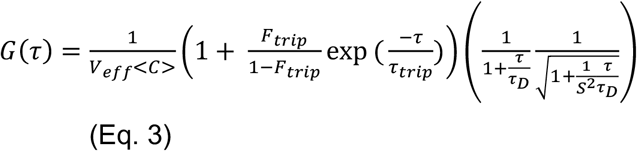

where C is the particle concentration, V_eff_ is the effective Gaussian volume of the focal spot described by ω_0_ (lateral radius) and z_o_ (axial radius). S is the structural parameter such that S = z_0_/ω_0_ and V_eff_ = π^3/2^ω_0_^2^z_0_. The mean number of molecules in the focus is given by *V*_*eff*_ < *C* >. F_trip_ is the fraction of triplet molecules and τ_trip_ is the relaxation time. τ_D_ is the characteristic dwell time required by molecules to diffuse through the focal volume from which the diffusion coefficient is calculated as follows:

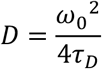

Since there exist two components of Fgf8-eGFP (N_1_/fast and N_2_/slow) [3], a two-component model was used to fit Fgf8-eGFP auto-correlation curves. The auto-correlation function from two species is the weighted sum of the individual correlation functions and is given by the following:

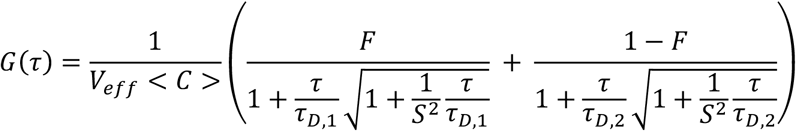

where *V*_*eff*_ < *C* >= *N*_*total*_ = *N*_1_+ *N*_2_, and F is the fraction of molecules of the first species and 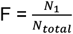 and 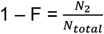

### Fluorescence cross-correlation (FCCS) analysis

FCCS was used to quantify the molecular interaction between Fgf8-eGFP and mRFP labeled extracellular HSPG [36, 37]. In this method, fluorescence intensity fluctuations are recorded individually from the green and red channel and the signal is then cross-correlated to obtain the cross-correlation function.

Fluorescence intensity in either channel is the sum of single labeled and double labeled molecules. The auto-correlation function for free-diffusion in 3D for total green and total red fluorescent molecules in eq. 3 (without triplet) is rewritten as:

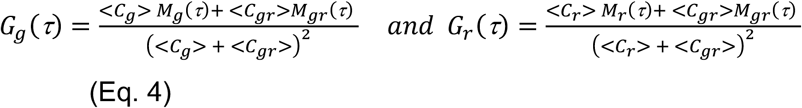

where

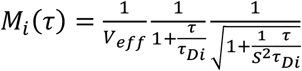

This is based on the simplest case where the effective detection volume is the same for both channels and overlaps perfectly. Cross-correlation function is then defined as follows:

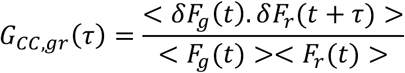

and is obtained from eq.4 as follows:

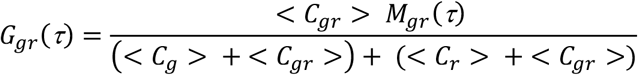

For quantification of interaction, the amplitude of auto-correlation and cross-correlation functions are used. Note that the amplitude of correlation function is related to the number of bound and unbound molecules (eq.2). At τ = 0 (maximum amplitude),

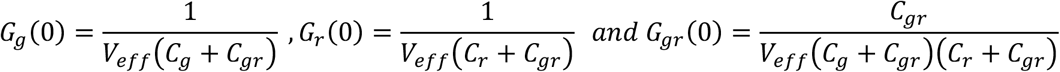

and a relative amplitude of the auto-correlation and cross-correlation functions then becomes:

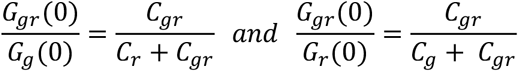

This relation gives the fraction of bound molecules relative to the red molecules or the green molecules and is denoted as the percentage of cross-correlation (% CCR).

### Evaluation of in vivo Dissociation constant K_d_

According to the law of mass action, the equilibrium dissociation constant is given by:

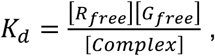

where [*R*_*free*_] is the concentration of free mRFP tagged HSPG proteins, [*G*_*free*_] for eGFP tagged ligand protein and [*Complex*] is the concentration of complex. The concentration of red, green and complex was calculated from the number of molecules in the focal volume obtained from auto and cross-correlation amplitudes.

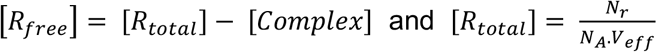

where N_A_ is the Avogadro’s number = 6.022 x 10^23^ mol^-1^. N_r_ is the number of molecules in the focal volume obtained from the auto-correlation amplitude in the red channel. Veff = π^3/2^ω_0_ z_0_, where ω_0_ = 0.214 μm and z_*0*_ = 1.19 μm. ω_0_ was determined by measuring the diffusion time of free dye CF568 in both channels and z_0_ was obtained from structure parameter (S) such that S = *z*_0_ /*ω*_0_. Similarly 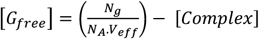 and 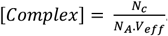.

K_d_ was then obtained from the slope of a plot between [*R*_*free*_]. [*G*_*free*_] and [*Complex*], following a linear regression fit.

### Fgf8 gradient measurements and nomenclature

For consistency, the text refers to the product of the zebrafish *fgf8a* gene [10], mutated in the *acerebellar* null mutant [11], as ‘Fgf8’; the duplicated paralogous *fgf8b* gene (previously *fgf17*), though similar in gain-of-function experiments, is not expressed at the gastrulation stages examined here [49, 50]. Embryos were injected at 1-cell stage with the desired mRNA or morpholino and at 32-cell stage with 20 pg of Fgf8-eGFP mRNA. For ectopic expression of Gpc4^ex^mRFP and Sdc2^ex^mRFP, 25 pg and 10 pg mRNA, respectively, was injected in 1-cell staged embryos. For knockdown of *gpc4* transcripts, 1 mM of morpholino was injected.

At the sphere stage, injected embryos were mounted in 1% agarose and imaged with Zeiss confocal LSM700 inverse microscope. Due to the injection in a single blastomere, a restricted source of cells was seen at the sphere stage and Fgf8-eGFP concentration decreased away from this source. Confocal image of a single plane was taken for multiple embryos, less than 20 μm inside the enveloping layer. Concentration of Fgf8-eGFP in the free extracellular space, was obtained by measuring the fluorescence intensity in the confocal images. Fluorescence intensity was measured inside a fixed circular space at different distances from the source, using ImageJ (Fiji) software.

The normalized concentration profile was modelled using the reaction diffusion equation for the particular geometry of the embryos and the result gave the decay length (*λ*) [3]. *λ* signifies the distance from the source at which Fgf8-eGFP concentration decays to approximately 50 % of its maximum and this was found to be around 200 μm or 10 cell-diameter.

### Membrane Scanning FCS

For auto/cross-correlation measurements in embryos, membrane scanning FCS (in Figure 4) was performed using the same Zeiss LSM780/Confocor 3 microscope and C-Apochromat 40x objective (N.A. 1.2), as mentioned above. The sample was excited using 488 nm line of Argon laser and 561 nm laser with laser powers ∼1 and ∼2 μW, respectively. A multi-track line scan mode was used to alternately excite the red and the green fluorophores. A confocal image of the membrane was first acquired. The direction of the line scan was then placed perpendicular to the membrane and a continuous line scan was started. The photon stream was spectrally split using a dichroic NFT 565 and emission filters LP580 and BP495-555, and then detected using APDs. The photon arrival times were recorded in the photon mode of the hardware correlator Flex 02-01D (correlator.com). Pinhole diameter of 40 μm was used (1 A.U for the red channel). Measurements were done typically for 300 s. Count rate for membrane bound Fgf8-eGFP was 8-20 kHz and for membrane bound mRFP-tagged proteins was 50-100 kHz. Data analysis was performed as described previously using a Matlab (Mathworks) program [16, 42]. Briefly, after binning of the photo stream in intervals of 2 μs, every line scan was arranged as a matrix. In each line, the position of the maximum corresponded to the membrane signal and membrane movements were corrected by aligning the maxima. An average over all rows was fitted with a Gaussian profile of width σ, and only the elements of each row between -2.5σ and 2.5σ were added to construct the intensity trace. From the resulting intensity trace over time, auto- and cross-correlation curves were computed with a multiple-tau correlation algorithm and fitted with a non-linear least squares fitting algorithm [42]. The number of molecules obtained after the fit were then used to evaluate the percentage of cross-correlation.

### Correlative Light Electron Microscopy (CLEM) of ultrathin cryo-sections

Embryos were fixed with 4% paraformaldehyde (PFA) in 0.1 M phosphate buffer (PB, pH 7.4) and processed for Tokuyasu cryo-sectioning as described [51, 52]. Embryos were washed in PB, infiltrated into 10% gelatin, cooled on ice, incubated in 2.3 M sucrose / water for 24 hours at 4°C, mounted on pins (Leica # 16701950), and plunge frozen in liquid nitrogen. 70 nm sections were cut on a Leica UC6+FC6 cyo-ultramicrotome and picked up in methyl cellulose / sucrose (1 part 2% methyl cellulose (MC), Sigma M-6385, 25 centipoises + 1 part 2.3 M sucrose).

To facilitate the identification of eGFP-labeled cells, sections were stained for CLEM [53]. For this, grids were placed upside down on drops of PBS in a 37°C-incubator for 20 min, washed with 0.1% glycin / PBS (5x 1 min), blocked with 1% BSA/PBS (2x 5 min) and incubated with rabbit anti-GFP (TP 401, Torrey Pines, 1:100) for 1 hour. After washes in PBS (4x 2 min), sections were incubated with Protein A conjugated to 10 nm gold for 1 hour, washed in PBS (3x 5 s, 4x 2 min) and post-fixed in 1% glutaraldehyde (5 min). After that sections were incubated with goat-anti-rabbit Alexa488 for the identification of Fgf8-eGFP positive cells in the fluorescence microscope, washed with PBS (4x 2 min), stained with 1 μg/ml DAPI for 10 min, and washed in water (10x 1 min). Grids were mounted in 50% glycerin/water between two coverslips and imaged at the Keyence Biozero 8000 fluorescence microscope. Sections were demounted, washed with distilled water (10x 1 min), stained with neutral uranyloxalate (2% uranylacetate (UA) in 0.15 M oxalic acid, pH 7.0) for 5 min, washed in water and incubated in MC containing 0.4% UA for 5 min. Grids were looped out, the MC/UA film was reduced to an even thin film and air dried. Finally, the sections were analyzed on a Morgagni 268 (FEI) at 80 kV and images were taken with a MegaView III digital camera (Olympus).

### Statistical Methods

Statistical analysis was performed using GraphPad Prism. The tests applied were unpaired Student’s T-test (assuming same variances in the two populations) and one-way ANOVA (assuming a Gaussian population distribution). Number of fish embryos used per experiment are mentioned in the text. FCS measurements were performed multiple times per embryo (>3) for multiple embryos (>15), for each test condition. An average of all measurements was used for statistical analysis. Data are represented as mean ± standard deviation (s.d.) or mean ± standard error of mean (s.e.m.). Statistical significance was defined as * P< 0.05, ** P< 0.01, *** P< 0.001, n.s. not significant.

### Heparan sulfate injections

Heparan sulfate injections were into the extracellular space of manually dechorionated 256-cell stage embryos. Dechorionation was conducted in petri dishes coated with 2 % agarose (in E3 buffer [45]) and the transfer of dechorionated embryos was done with glass pasteur pipettes to avoid damage. For injections, embryos were transferred onto agarose plates containing cubical depressions, for stabilization. Each embryo was injected with 2 nl of HS mix (90% HS (0,5 ng/ml; Sigma Aldrich H7640) + 10% phenol red (PR)) or CTRL mix (90% nuclease free water + 10% PR), respectively. The needles were loaded with solution and mounted onto a micromanipulation device (MM33, WPI). Prior to the usage of each needle the injection volume was calibrated by injecting the respective solution into an oil droplet and measuring its diameter with the ocular scale. Measurements of expression ranges of Fgf target genes in HS injected embryos after *in-situ* hybridisation (ISH) were performed with the line measurement tool in Fiji. In this experiment the different groups were blinded by a colleague to avoid bias. Statistical significance of differences between the investigated groups was determined by calculating the p-values using GraphPad Prism.

## Supplemental Figure Legends

**Figure S1: Fgf signaling is upregulated by ectopic expression of extracellular and not cell-surface attached HSPG**.

**(a,b)** Illustration of constructs used for ectopic expression of Gpc and Sdc proteins. **(c-f)** Expression pattern of Fgf downstream targets *spry2, spry4* and *etv4* during gastrulation (60% epiboly) in wild-type (Wt) compared to injected embryos, tested by *in-situ* hybridization: lateral views with animal pole to the top, dorsal to the right. 100 pg mRNA was injected for all indicated Gpc and Sdc at one-cell stage. **(c-d)** Ectopic expression of 10 different full-length Gpc (Gpc-fl, c) or 3 different Sdc family members (Sdc-fl,d) did not influence Fgf downstream target expression. **(e-f)** Ectopic expression of extracellular-mRFP tagged Gpc and Sdc (named Gpc^ex^mRFP and Sdc^ex^mRFP). **(e)** After mRNA injection of extracellular domain of several Gpc, only Gpc1b^ex^mRFP, Gpc4^ex^mRFP and Gpc6a^ex^mRFP resulted in an expanded expression pattern of *spry2* and *etv4*. **(f)** Ectopic expression of Sdc2^ex^mRFP, Sdc3^ex^mRFP and Sdc4^ex^mRFP also increased *spry2* and *etv4* expression pattern. Gpc2^ex^mRFP and Gpc5c^ex^mRFP could not be tested as the resulting fusion proteins were not localized extracellularly (Figure S2p,q). **(g)** Fgf8 mRNA expression domain was not significantly affected after injecting 100 pg each of extracellular Gpc (Gpc1b^ex^mRFP, Gpc4^ex^mRFP and Gpc6a^ex^mRFP) and Sdc (Sdc2^ex^mRFP, Sdc3^ex^mRFP and Sdc4^ex^mRFP). **(h,i)** Heparan sulfate (HS) injection in zebrafish embryos causes dorsalization and alters Fgf target gene expression. **(h)** Dorsalized phenotype caused by HS injection. Embryos were injected with either HS or CTRL solution (90% nuclease free water + 10% PR) at the 256-cell stage. For quantification, yolk dimensions were measured for each group at the tailbud stage (red double arrow line) using the line measurement function in Fiji, placed between the polster and the tailbud. Scale bar = 250 μm. Quantification of the elongation phenotype. One-way ANOVA, Bonferroni *post-hoc* test,** p < 0.01, **** p< 0.0001. Uninjected n = 28, CTRL injected n = 32, HS injected n = 20. **(i)** HS injection alters Fgf target gene expression in zebrafish embryos. Embryos were injected at the 256-cell stage with HS or CTRL solution and fixed at the 70-80 % epiboly stage for ISH. The width of the *spry4* expression domain at the embryonic margin was measured for each group along the red bracket, using the line measurement tool in Fiji. Scale bar = 250 μm. Quantification of the *spry4* expression range at the margin. One-way ANOVA, Bonferroni *post-hoc* test,** p < 0.01, **** p< 0.0001. fl: full length, ss: signal sequence, Gpc: glypican domain, Sdc-ex: syndecan extracellular domain, GPI: GPI anchor, Sdc-tm: syndecan transmembrane domain, cy: cytoplasmic domain. Scale bar: 200 μm.

**Figure S2: Localization of different fluorescent-tagged HSPG fusion proteins**.

(a-d) mRFP or eGFP was fused with different HSPGs at positions along the protein length such that the membrane anchor remains unaffected. mRNA from the resulting fusion proteins was injected at one-cell stage and imaged at 5 hpf. Shown here are examples of each HSPG family. (a-c) mRFP or eGFP tagged at the N-terminal, between the signal sequence and protein domain of Gpc4 (a), Gpc3 (b) or Sdc2 (c) resulted in fluorescence on the cell membrane (hollow arrow) and ECM (filled arrow), implying protein release from the cell membrane. (d) Disruption of the Gpc4 protein by placing mRFP in between Gpc domain and GPI anchor, abolished cleavage from the membrane (hollow arrow). ss: signal sequence, GPI: GPI anchor, tm: transmembrane domain, cy: cytoplasmic domain. Scale bar: 10 μm. (e-q) Localization of extracellular mRFP tagged HSPGs as illustrated in Figure S1a-b. (e-o) Most mRFP-tagged HSPG constructs were found secreted in the extracellular space. (p-q) Gpc2^ex^mRFP (p) and Gpc5c^ex^mRFP (q) were not secreted (arrows) and localized inside the cells, hence could not be used for FCCS. Different constructs with varied linkers were tested, but every time, Gpc2^ex^mRFP and Gpc5c^ex^mRFP were intracellular (not shown). Scale bar: 20 μm.

**Figure S3: K**_**d**_ **analysis of Fgf8 and HSPG binding in the extracellular space**.

Scatter plot for evaluation of effective dissociation constant (K_d_) for binding between Fgf8-eGFP and various HSPG: (a) glypican family 1/2/4/6, (b) glypican family 3/5 and (c) syndecans. The graph represents a product of concentrations of free red ([R_free_]) and green ([G_free_]) molecules *versus* the concentration of complex ([Complex]). Concentration values were obtained from FCCS measurements. Solid line shows the linear fit of the data. Dissociation constant values K_d_ are mentioned in red for each HSPG.

**Figure S4: Removal of HS side chains abolishes interaction of Fgf8 with extracellular glypicans and syndecans**.

(a) Point mutation to remove HS attachment sites from glypicans and syndecans. HS chains are covalently attached to Serine-Glycine (SG) residues (blue) on the core protein. There are 2-4 SG repeats in glypicans present more of less contiguously, but in syndecans, there are multiple copies (3-6), spanning the entire N-terminal domain (Sdc2, Sdc4) or even the entire gene (Sdc3) [47]. All the serine residues from SG pairs were mutated to alanine (A) to generate Gpc^ex^mRFP-HS and Sdc^ex^mRFP-HS. The resulting constructs were also localized to the extracellular space after mRNA injection (not shown). ss: signal sequence, Gpc: glypican domain, Sdc: syndecan domain. (b) Percentage of cross-correlation for Fgf8-eGFP and HSPG with and without HS side chains. Gpc1b^ex^mRFP, Gpc4^ex^mRFP, Gpc6a^ex^mRFP, Sdc2^ex^mRFP, Sdc3^ex^mRFP and Sdc4^ex^mRFP which influenced Fgf8 downstream signaling (Figure S1e,f), were tested for binding to Fgf8-eGFP in the absence of their HS side chains. Note the loss in cross-correlation upon removal of HS chains. The % CCR after loss of HS chains was not significant compared to the negative control (sec-eGFP vs sec-mRFP). Graph represents mean with s.d. (c) Slow fraction of Fgf8 is not affected after ectopic expression of Gpc4^ex^mRFP and Sdc2^ex^mRFP in embryos. Percentage of slow fraction was determined using a 2-component fit to FCS autocorrelation data from Figure 2. Graph represents mean with s.d. Data was analyzed using one-way ANOVA. (d) Injection of morpholinos against Gpc4 recapitulated the shortened tail phenotype obtained in *gpc4* mutant [41]. Wild-type embryos at 24 hpf are compared to embryos injected with 1 mM of translational block morpholino (Gpc4-MO1) and 1 mM of splice-junction morpholino (Gpc-MO2). The embryos are also delayed in development. Scale bar: 200 μm. Gpc4-MO1 was used for further experiments. (e) Upon injection of Gpc4^ex^mRFP mRNA, fluorescence can be seen at the animal pole of sphere stage embryos. Co-injection of Gpc4^ex^mRFP mRNA with 1 mM of Gpc4-MO1 resulted in the loss of fluorescent signal, indicating that *gpc4* mRNA was degraded. Scale bar: 500 μm. Brightfield images are also shown. (f) Concentration profile of Fgf8-eGFP becomes steeper upon over-expression of cell-surface attached Gpc4. Fgf8-eGFP gradient in wild-type embryos (Wt) is compared to embryos injected with different mRNA concentrations of cell-membrane attached Gpc4 (mRFP-Gpc4) (50 pg; n= 21 and 200 pg, n=17). Steeper gradient is formed due to the release of Gpc4 into the ECM, which binds and prevents Fgf8 from further diffusion. (g) Plot of decay length (*λ*) for the different conditions, showing a reduction in decay length. Statistical significance was inferred using One-way ANOVA. Error bars: s.d.

**Figure S5: Correlative light and electron microscopy to study the extracellular distribution of Fgf8-eGFP molecules around cell membranes**.

(a-d) Examples from 4 different embryos are shown, highlighting the trapping of Fgf8 molecules in the extracellular space. 70 nm thick sections from embryos injected with Fgf8-eGFP were immunostained with rabbit anti-GFP protein A gold, goat anti-rabbit Alexa488 and DAPI. Regions with strong fluorescence (arrows and boxes) were further imaged with the electron microscope and to detect labeling with 10 nm gold particles (Materials and Methods). All examples highlight regions which are sandwiched by two cell membranes. Gold labeled particles can be seen on the plasma membrane (pm) as well as attached to a grayish mesh in the ECM. Matrix associated Fgf8 molecules are distributed up to 50 nm away from the membrane. Interestingly Fgf8 molecules were found excluded from membrane protrusions (b). Arrowheads in (d) also depict Fgf8-eGFP.

## References

1. Wolpert, L., 1969. Positional information and the spatial pattern of cellular differentiation. Journal of Theoretical Biology. 25(1): p. 1–47. DOI: 10.1016/S0022-5193(69)80016-0.

2. Rogers, K.W. and Schier, A.F., 2011. Morphogen Gradients: From Generation to Interpretation. Annual Review of Cell and Developmental Biology. 27(1): p. 377–407. DOI: 10.1146/annurev-cellbio-092910-154148.

3. Yu, S.R., Burkhardt, M., Nowak, M., Ries, J., Petrášek, Z., Scholpp, S., Schwille, P., and Brand, M., 2009. Fgf8 morphogen gradient forms by a source-sink mechanism with freely diffusing molecules. Nature. 461(7263): p. 533–36. DOI: 10.1038/nature08391.

4. Wang, Y., Wang, X., Wohland, T., and Sampath, K., 2016. Extracellular interactions and ligand degradation shape the nodal morphogen gradient. Elife. 5. DOI: 10.7554/eLife.13879.

5. Stapornwongkul, K.S. and Vincent, J.P., 2021. Generation of extracellular morphogen gradients: the case for diffusion. Nat Rev Genet. 22(6): p. 393–411. DOI: 10.1038/s41576-021-00342-y.

6. Lord, N.D., Carte, A.N., Abitua, P.B., and Schier, A.F., 2021. The pattern of nodal morphogen signaling is shaped by co-receptor expression. Elife. 10. DOI: 10.7554/eLife.54894.

7. Harish, R.K., Gupta, M., Zoller, D., Hartmann, H., Gheisari, A., Machate, A., Hans, S., and Brand, M., 2023. Real-time monitoring of an endogenous Fgf8a gradient attests to its role as a morphogen during zebrafish gastrulation. Development. 150(19). DOI: 10.1242/dev.201559.

8. Cohn, M.J., Izpisua-Belmonte, J.C., Abud, H., Heath, J.K., and Tickle, C., 1995. Fibroblast growth factors induce additional limb development from the flank of chick embryos. Cell. 80(5): p. 739–46. DOI: 10.1016/0092-8674(95)90352-6.

9. Crossley, P.H., Martinez, S., and Martin, G.R., 1996. Midbrain development induced by FGF8 in the chick embryo. Nature. 380(6569): p. 66–8. DOI: 10.1038/380066a0.

10. Reifers, F., Böhli, H., Walsh, E.C., Crossley, P.H., Stainier, D.Y.R., and Brand, M., 1998. Fgf8 is mutated in zebrafish acerebellar (ace) mutants and is required for maintenance of midbrain-hindbrain boundary development and somitogenesisy. Development. 125(13): p. 2381–95. DOI: 10.1242/dev.125.13.2381.

11. Brand, M., Heisenberg, C.P., Jiang, Y.J., Beuchle, D., Lun, K., Furutani-Seiki, M., Granato, M., Haffter, P., Hammerschmidt, M., Kane, D.A., et al., 1996. Mutations in zebrafish genes affecting the formation of the boundary between midbrain and hindbrain. Development. 123: p. 179–90. DOI: 10.1242/dev.123.1.179.

12. Bökel, C. and Brand, M., 2013. Generation and interpretation of FGF morphogen gradients in vertebrates. Curr Opin Genet Dev. 23(4): p. 415–22. DOI: 10.1016/j.gde.2013.03.002.

13. Bökel, C. and Brand, M., 2014. Endocytosis and signaling during development. Cold Spring Harb Perspect Biol. 6(3). DOI: 10.1101/cshperspect.a017020.

14. Zhou, S., Lo, W.-C., Suhalim Jeffrey L., Digman Michelle A., Gratton, E., Nie, Q., and Lander Arthur D., 2012. Free Extracellular Diffusion Creates the Dpp Morphogen Gradient of the Drosophila Wing Disc. Current Biology. 22(8): p. 668–75. DOI: 10.1016/j.cub.2012.02.065.

15. Müller, P., Rogers, K.W., Jordan, B.M., Lee, J.S., Robson, D., Ramanathan, S., and Schier, A.F., 2012. Differential Diffusivity of Nodal and Lefty Underlies a Reaction-Diffusion Patterning System. Science. 336(6082): p. 721–24. DOI: 10.1126/science.1221920.

16. Ries, J., Yu, S.R., Burkhardt, M., Brand, M., and Schwille, P., 2009. Modular scanning FCS quantifies receptor-ligand interactions in living multicellular organisms. Nature Methods. 6(9): p. 643–45. DOI: 10.1038/nmeth.1355.

17. Kicheva, A., Bollenbach, T., Wartlick, O., Julicher, F., and Gonzalez-Gaitan, M., 2012. Investigating the principles of morphogen gradient formation: from tissues to cells. Curr Opin Genet Dev. 22(6): p. 527–32. DOI: 10.1016/j.gde.2012.08.004.

18. Ng, X.W., Sampath, K., and Wohland, T., 2018. Fluorescence Correlation and Cross-Correlation Spectroscopy in Zebrafish. Methods Mol Biol. 1863: p. 67–105. DOI: 10.1007/978-1-4939-8772-6_5.

19. Matsuda, S. and Affolter, M., 2023. Is Drosophila Dpp/BMP morphogen spreading required for wing patterning and growth? Bioessays. 45(9): p. e2200218. DOI: 10.1002/bies.202200218.

20. Duchesne, L., Octeau, V., Bearon, R.N., Beckett, A., Prior, I.A., Lounis, B., and Fernig, D.G., 2012. Transport of fibroblast growth factor 2 in the pericellular matrix is controlled by the spatial distribution of its binding sites in heparan sulfate. PLoS Biol. 10(7): p. e1001361. DOI: 10.1371/journal.pbio.1001361.

21. Yan, D. and Lin, X., 2009. Shaping Morphogen Gradients by Proteoglycans. Cold Spring Harbor Perspectives in Biology. 1(3): p. a002493–a93. DOI: 10.1101/cshperspect.a002493.

22. Matsuo, I. and Kimura-Yoshida, C., 2013. Extracellular modulation of Fibroblast Growth Factor signaling through heparan sulfate proteoglycans in mammalian development. Current Opinion in Genetics & Development. 23(4): p. 399–407. DOI: 10.1016/j.gde.2013.02.004.

23. Balasubramanian, R. and Zhang, X., 2016. Mechanisms of FGF gradient formation during embryogenesis. Seminars in Cell & Developmental Biology. 53: p. 94–100. DOI: 10.1016/j.semcdb.2015.10.004.

24. Chan, W.K., Price, D.J., and Pratt, T., 2017. FGF8 morphogen gradients are differentially regulated by heparan sulphotransferases Hs2st and Hs6st1 in the developing brain. Biol Open. 6(12): p. 1933–42. DOI: 10.1242/bio.028605.

25. Lin, X., 2004. Functions of heparan sulfate proteoglycans in cell signaling during development. Development. 131(24): p. 6009–21. DOI: 10.1242/dev.01522.

26. Choi, Y., Chung, H., Jung, H., Couchman, J.R., and Oh, E.-S., 2011. Syndecans as cell surface receptors: Unique structure equates with functional diversity. Matrix Biology. 30(2): p. 93–99. DOI: 10.1016/j.matbio.2010.10.006.

27. Gupta, M. and Brand, M., 2013. Identification and expression analysis of zebrafish glypicans during embryonic development. PLoS One. 8(11): p. e80824. DOI: 10.1371/journal.pone.0080824.

28. Manon-Jensen, T., Itoh, Y., and Couchman, J.R., 2010. Proteoglycans in health and disease: the multiple roles of syndecan shedding. The FEBS Journal. 277(19): p. 3876–89. DOI: 10.1111/j.1742-4658.2010.07798.x.

29. Traister, A., Shi, W., and Filmus, J., 2008. Mammalian Notum induces the release of glypicans and other GPI-anchored proteins from the cell surface. Biochemical Journal. 410(3): p. 503–11. DOI: 10.1042/BJ20070511.

30. Capurro, M.I., Shi, W., and Filmus, J., 2012. LRP1 mediates Hedgehog-induced endocytosis of the GPC3–Hedgehog complex. Development. 139(20): p. e1–e1. DOI: 10.1242/dev.089300.

31. Capurro, M., Martin, T., Shi, W., and Filmus, J., 2014. Glypican-3 binds to frizzled and plays a direct role in the stimulation of canonical Wnt signaling. Journal of Cell Science. p. jcs.140871. DOI: 10.1242/jcs.140871.

32. Scholpp, S. and Brand, M., 2004. Endocytosis Controls Spreading and Effective Signaling Range of Fgf8 Protein. Current Biology. 14(20): p. 1834–41. DOI: 10.1016/j.cub.2004.09.084.

33. Fürthauer, M., Van Celst, J., Thisse, C., and Thisse, B., 2004. Fgf signalling controls the dorsoventral patterning of the zebrafish embryo. Development. 131(12): p. 2853–64. DOI: 10.1242/dev.01156.

34. Fürthauer, M., Reifers, F., Brand, M., Thisse, B., and Thisse, C., 2001. sprouty4 acts in vivo as a feedback-induced antagonist of FGF signaling in zebrafish. Development. 128(12): p. 2175–86. DOI: 10.1242/dev.128.12.2175.

35. Lambaerts, K., Van Dyck, S., Mortier, E., Ivarsson, Y., Degeest, G., Luyten, A., Vermeiren, E., Peers, B., David, G., and Zimmermann, P., 2012. Syntenin, a syndecan adaptor and an Arf6 phosphatidylinositol 4,5-bisphosphate effector, is essential for epiboly and gastrulation cell movements in zebrafish. Journal of Cell Science. 125(5): p. 1129–40. DOI: 10.1242/jcs.089987.

36. Bacia, K., Kim, S.A., and Schwille, P., 2006. Fluorescence cross-correlation spectroscopy in living cells. Nature Methods. 3(2): p. 83–89. DOI: 10.1038/nmeth822.

37. Bacia, K. and Schwille, P., 2007. Practical guidelines for dual-color fluorescence cross-correlation spectroscopy. Nature Protocols. 2(11): p. 2842–56. DOI: 10.1038/nprot.2007.410.

38. Müller, P., Rogers, K.W., Yu, S.R., Brand, M., and Schier, A.F., 2013. Morphogen transport. Development. 140(8): p. 1621–38. DOI: 10.1242/dev.083519.

39. Shimokawa, K., Kimura-Yoshida, C., Nagai, N., Mukai, K., Matsubara, K., Watanabe, H., Matsuda, Y., Mochida, K., and Matsuo, I., 2011. Cell Surface Heparan Sulfate Chains Regulate Local Reception of FGF Signaling in the Mouse Embryo. Developmental Cell. 21(2): p. 257–72. DOI: 10.1016/j.devcel.2011.06.027.

40. Kramer, K.L. and Yost, H.J., 2003. Heparan Sulfate Core Proteins in Cell-Cell Signaling. Annual Review of Genetics. 37(1): p. 461–84. DOI: 10.1146/annurev.genet.37.061103.090226.

41. Topczewski, J., Sepich, D.S., Myers, D.C., Walker, C., Amores, A., Lele, Z., Hammerschmidt, M., Postlethwait, J., and Solnica-Krezel, L., 2001. The Zebrafish Glypican Knypek Controls Cell Polarity during Gastrulation Movements of Convergent Extension. Developmental Cell. 1(2): p. 251–64. DOI: 10.1016/S1534-5807(01)00005-3.

42. Ries, J. and Schwille, P., 2006. Studying Slow Membrane Dynamics with Continuous Wave Scanning Fluorescence Correlation Spectroscopy. Biophysical Journal. 91(5): p. 1915–24. DOI: 10.1529/biophysj.106.082297.

43. Mohammadi, M., Olsen, S.K., and Goetz, R., 2005. A protein canyon in the FGF-FGF receptor dimer selects from an a la carte menu of heparan sulfate motifs. Curr Opin Struct Biol. 15(5): p. 506–16. DOI: 10.1016/j.sbi.2005.09.002.

44. Simon, N., Safyan, A., Pyrowolakis, G., and Matsuda, S., 2023. Dally is not essential for Dpp spreading or internalization but for Dpp stability by antagonizing Tkv-mediated Dpp internalization. eLife. DOI: 10.7554/eLife.86663.1.

45. Brand, M., Granato, M., and Nüsslein-Volhard, C., Keeping and raising zebrafish, in Zebrafish: A Practical Approach, C.N.-V.a.R. Dahm, Editor. 2002, Oxford University Press.: Oxford.

46. Kimmel, C.B., Ballard, W.W., Kimmel, S.R., Ullmann, B., and Schilling, T.F., 1995. Stages of embryonic development of the zebrafish. Developmental Dynamics. 203(3): p. 253–310. DOI: 10.1002/aja.1002030302.

47. Zhang, L., David, G., and Esko, J.D., 1995. Repetitive Ser-Gly Sequences Enhance Heparan Sulfate Assembly in Proteoglycans. Journal of Biological Chemistry. 270(45): p. 27127–35. DOI: 10.1074/jbc.270.45.27127.

48. Petrášek, Z. and Schwille, P., 2008. Precise Measurement of Diffusion Coefficients using Scanning Fluorescence Correlation Spectroscopy. Biophysical Journal. 94(4): p. 1437–48. DOI: 10.1529/biophysj.107.108811.

49. Reifers, F., Adams, J., Mason, I., Schulte-Merker, S., and Brand, M., 2000. Overlapping and distinct functions provided by fgf17, a new zebrafish member of the Fgf8/17/18 subgroup of Fgfs. Mechanisms of Development. 99: p. 39–49.

50. Jovelin, R., Yan, Y.L., He, X., Catchen, J., Amores, A., Canestro, C., Yokoi, H., and Postlethwait, J.H., 2010. Evolution of developmental regulation in the vertebrate FgfD subfamily. J Exp Zool B Mol Dev Evol. 314(1): p. 33–56. DOI: 10.1002/jez.b.21307.

51. Slot, J.W. and Geuze, H.J., 2007. Cryosectioning and immunolabeling. Nature Protocols. 2(10): p. 2480–91. DOI: 10.1038/nprot.2007.365.

52. Tokuyasu, K.T., 1980. Immunochemistry on ultrathin frozen sections. The Histochemical Journal. 12(4): p. 381–403. DOI: 10.1007/BF01011956.

53. Fabig, G., Kretschmar, S., Weiche, S., Eberle, D., Ader, M., and Kurth, T., Labeling of Ultrathin Resin Sections for Correlative Light and Electron Microscopy, in Methods in Cell Biology. 2012, Elsevier. p. 75–93.

