## Supplemental Figures for "Fine-tuning of Fgf8 morphogen gradient by heparan sulfate proteoglycans in the extracellular matrix"

### Supplementary Figure 1

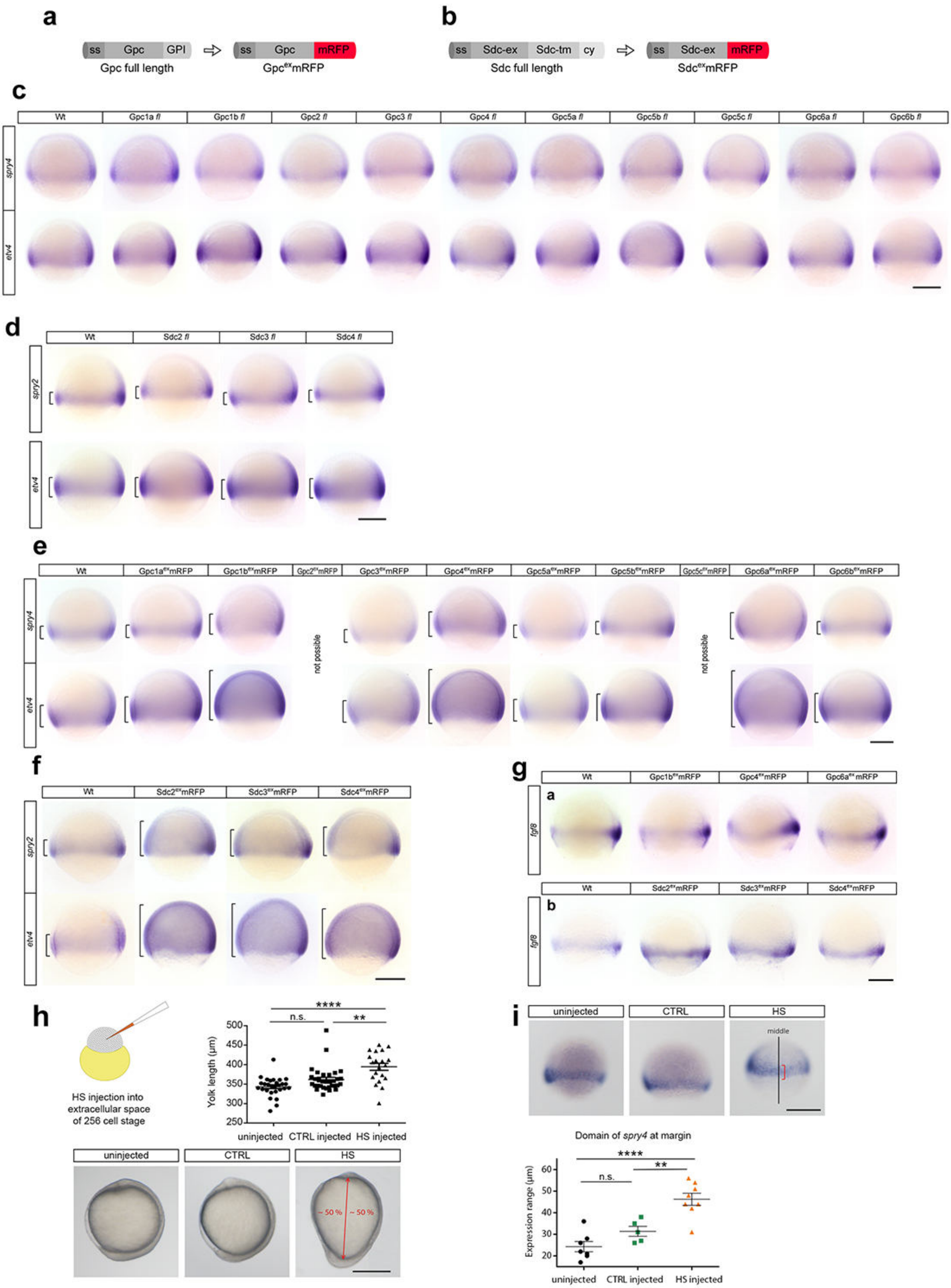

Supplementary Figure 2

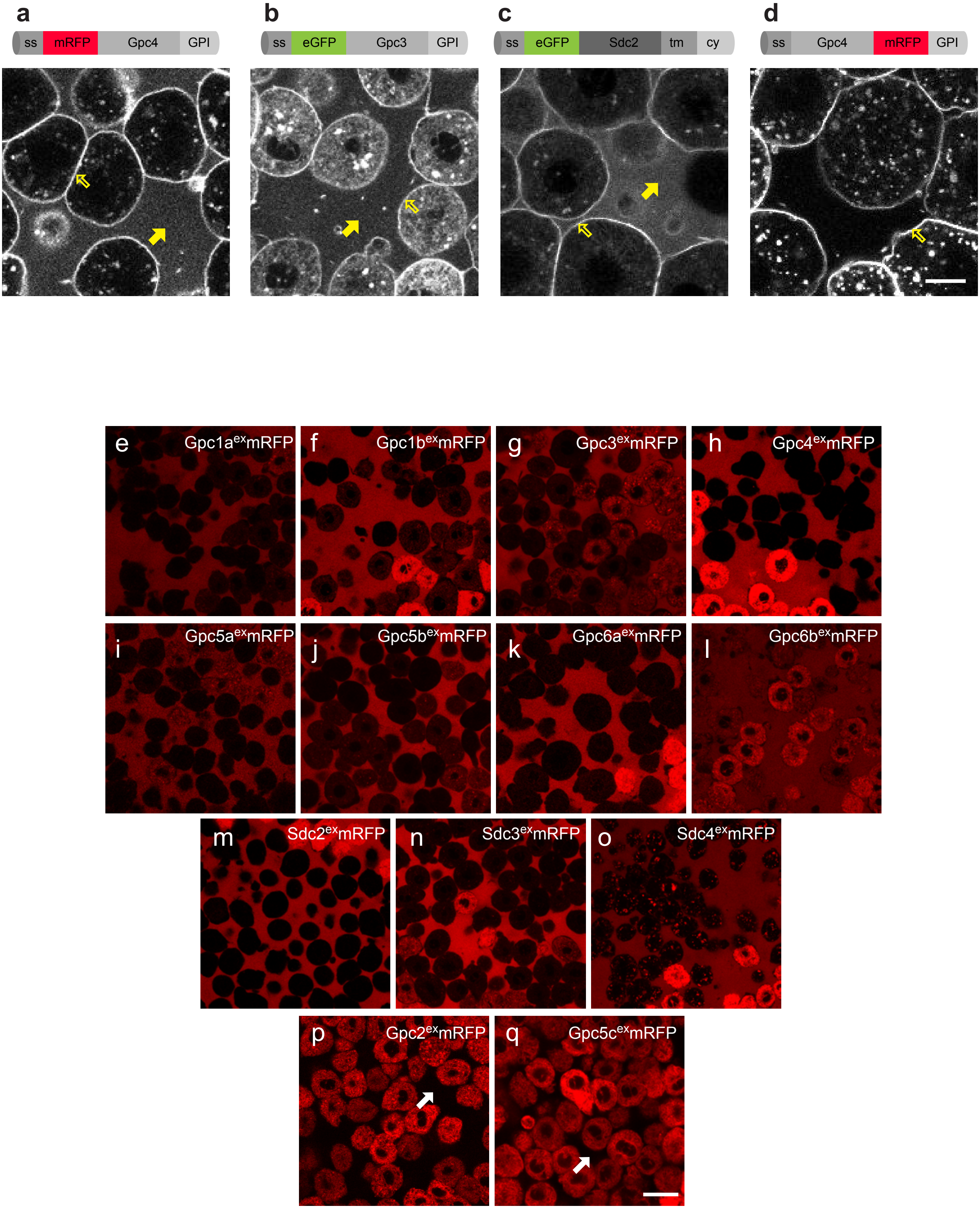

### Supplementary Figure 3

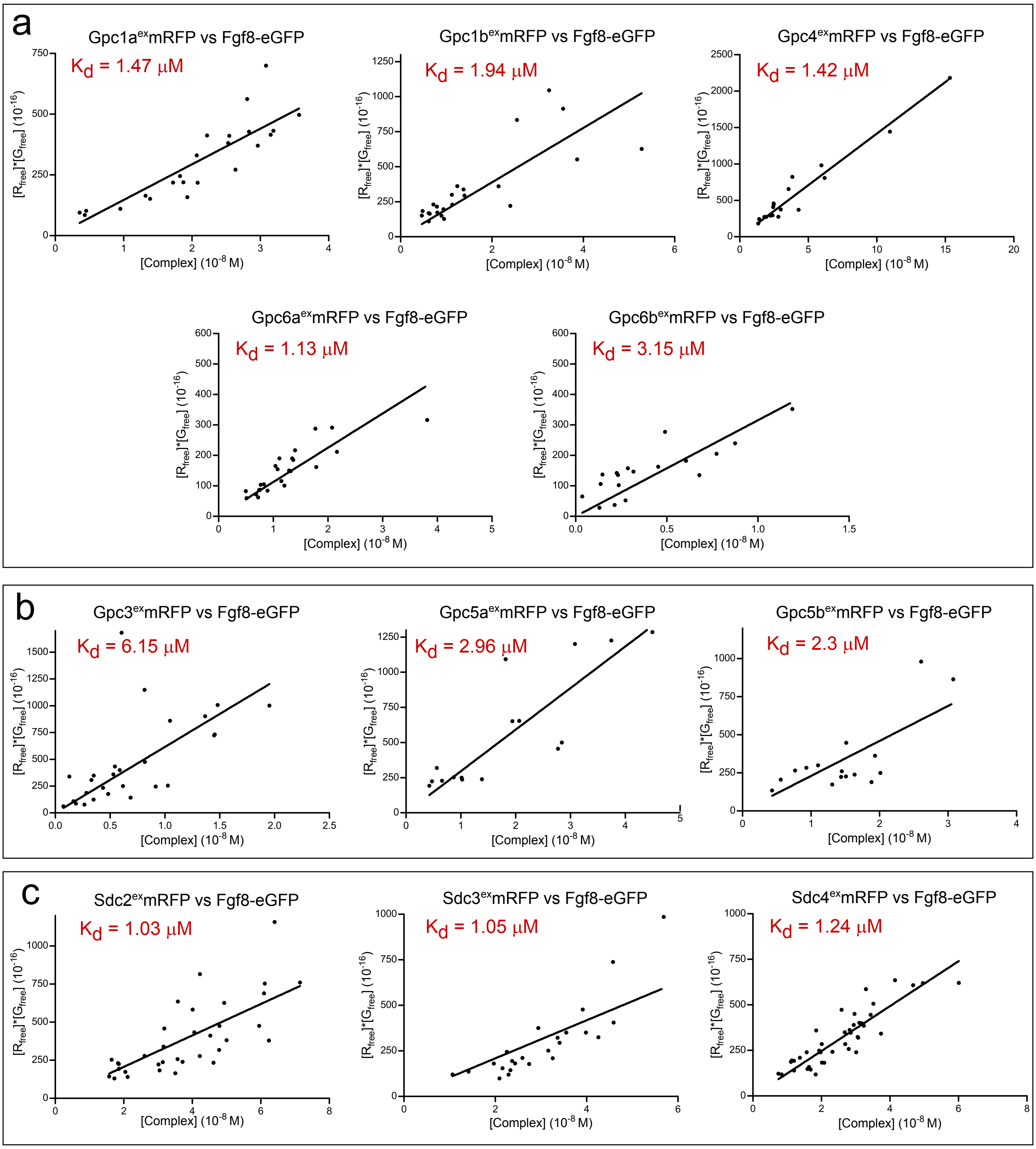

### Supplementary Figure 4

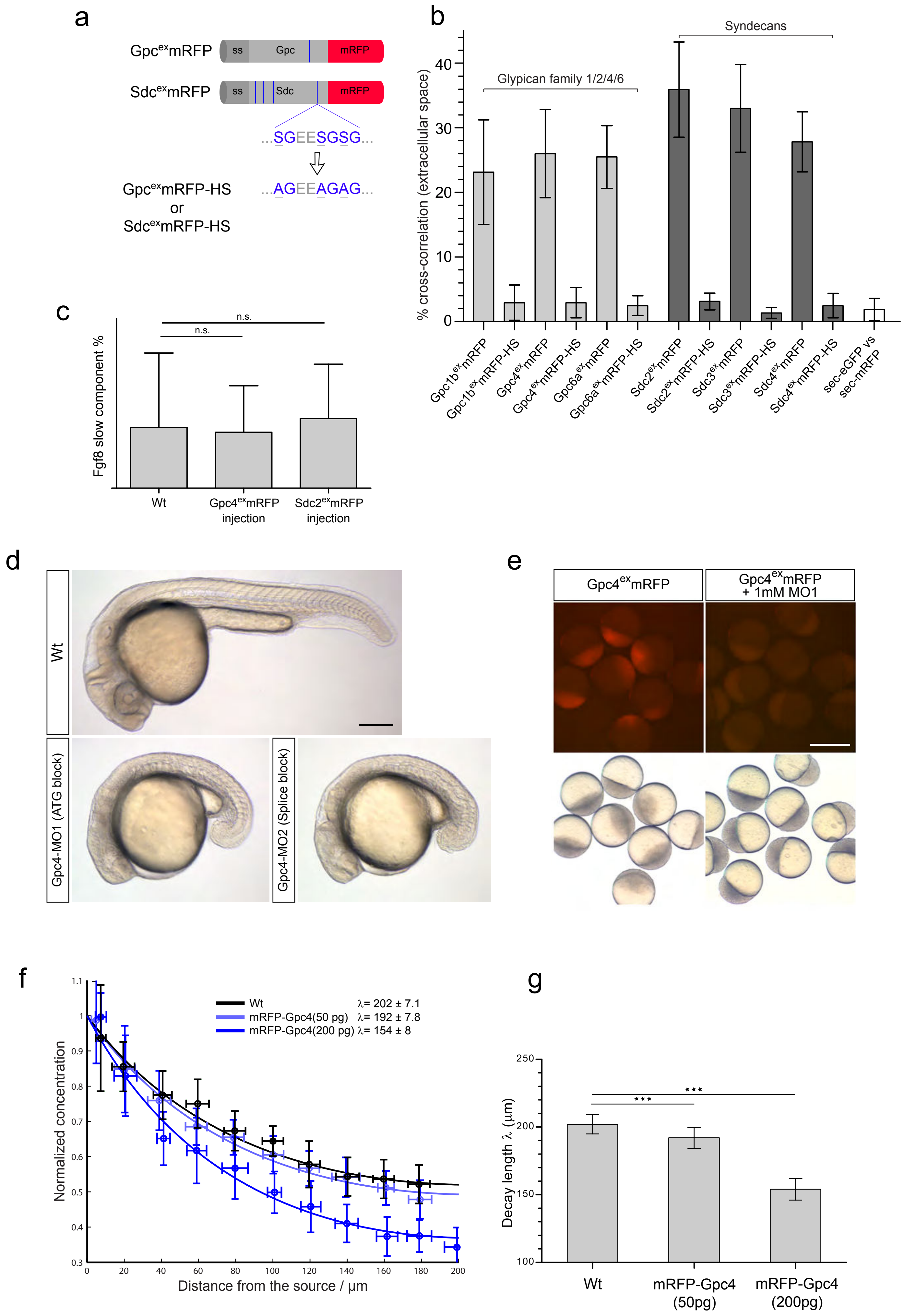

Supplementary Figure 5

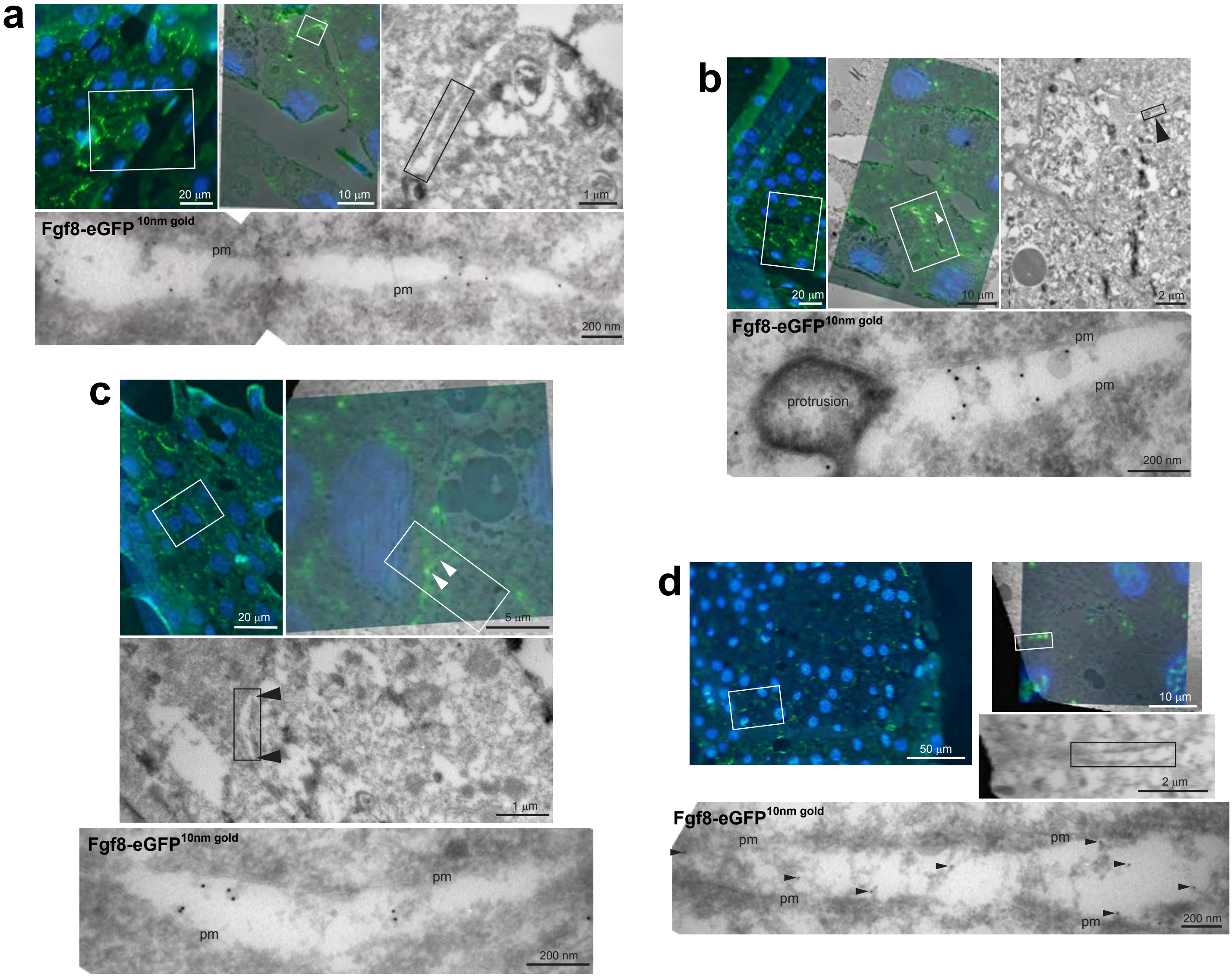
